# Histone deacetylase activity limits the response to EZH2 inhibition-based therapy in epithelioid sarcoma and is targetable by epigenetic combination

**DOI:** 10.64898/2026.08.04.742011

**Authors:** Noemi Arrighetti, Cesare Soffientini, Valentina Zuco, Stefano Percio, Loredana Cleris, Suzan Abdulrazak Ahmed, Elisa Del Savio, Luca Sigalotti, Roberta Maestro, Silvia Brich, Gian Paolo Dagrada, Marta Barisella, Paola Collini, Alex Kentsis, Paul H. Huang, Alessandro Gronchi, Anna Maria Frezza, Silvia Stacchiotti, Nadia Zaffaroni, Sandro Pasquali

## Abstract

Epithelioid sarcoma (EpS) is an ultra-rare, aggressive soft tissue sarcoma (STS) driven by INI1 loss and consequent hyperactivation of the chromatin-modifying enzyme EZH2. Although the EZH2 inhibitor tazemetostat has shown clinical activity, responses remain limited, highlighting the need for improved treatment strategies. Here, using two in-house generated patient-derived xenograft models and matched cell lines derived from INI-1 deficient EpS, we investigated EZH2 inhibition in combination with doxorubicin, the first-line standard for advanced STSs, identifying distinct patterns of response and resistance. Integrated transcriptomic and functional analyses revealed that response to EZH2 inhibition-based therapy was associated with chromatin remodeling, characterized by increased H3K27 acetylation and downregulation of histone deacetylase (HDAC)-related transcriptional programs. Conversely, the intrinsically resistant model failed to undergo this epigenetic transition despite EZH2 inhibition. Pharmacological HDAC inhibition restored H3K27 acetylation, promoted apoptosis, and enhanced the activity of EZH2 inhibition-based therapy. These findings identify failure to accumulate H3K27 acetylation as a hallmark of resistance to EZH2 inhibition-based treatment, and show that pharmacological HDAC inhibition can restore this chromatin transition and re-sensitize resistant tumors, providing a rationale for combined epigenetic targeting strategies in INI1-deficient malignancies.

**Translational relevance:** Prospective trials are challenging in rare tumors such as epithelioid sarcoma (EpS), limiting the level of evidence for existing therapies and the development of new agents. This is particularly relevant for EpS, where drug regimens are those used for all soft tissue sarcomas (STSs), and the specific mechanisms of drug response remain poorly understood. This preclinical study of tazemetostat in combination with doxorubicin shows differential outcomes in two INI1-deficient proximal-type EpS models, providing evidence of the heterogeneity that exists even within the same tumor subtype and fostering the need to better understand the molecular mechanisms driving drug sensitivity/resistance in this disease. In addition, we demonstrated the potential to treat EpS models through modulation of epigenetic mechanisms, showing that HDAC inhibition may restore sensitivity to EZH2-targeted therapy. These findings support the rationale for developing combination strategies incorporating epigenetic modulators and provide a preclinical framework for overcoming resistance to current therapies in EpS.

## Introduction

Epithelioid sarcoma (EpS) is an ultra-rare, aggressive soft-tissue sarcoma (STS) (1–5), characterized by a high tendency toward local recurrence and metastatic spread and associated with a dismal survival for patients with advanced disease (6). Two histologic subtypes are recognized on a morphological basis: the classical (or “distal”-type) (D-EpS), usually but not exclusively developing in distal extremities, and the proximal-type (P-EpS), more often arising in proximal extremities, perineum and trunk (7–11). Morphological differences are paralleled by distinct transcriptional and DNA methylation profiles, with D-EpS exhibiting an endothelial-like molecular makeup, whereas P-EpS displaying hyperactivation of MYC and GATA3 signaling pathways, resulting in worse clinical outcomes (7–13). The molecular hallmark of all EpS cases is the loss of expression of INI1, encoded by the *SMARCB1* gene. INI1 is a core component of the SWI/SNF chromatin remodeling complex and its loss abrogates SWI/SNF function and confers dependence on the opposing polycomb repressive complex 2 (PRC2), driven by the H3K27 methyltransferase EZH2 (14–17). EZH2 activity enforces a repressive chromatin state that silences differentiation and cell-cycle control programs, representing both a central pathogenic axis and a tractable therapeutic vulnerability in INI1-deficient tumors including EpS (18,19). Conversely, acetylation of H3K27 (H3K27ac) marks transcriptionally active chromatin and is mutually exclusive with H3K27me^3^ at the same residue. The dynamic balance between these two modifications thus represents a key regulatory switch governing gene expression at PRC2 loci (20). For patients with advanced disease, treatment options are limited (3,21). Anthracycline-based chemotherapy, which is the standard first-line treatment for STS, can be active in EpS, especially in the proximal subtype, but it is often associated with short-lived responses (21). The first-in-class EZH2 inhibitor (EZH2i) EPZ-6438 (tazemetostat, TZM) gained FDA approval in 2020 for advanced EpS following results of a phase II trial reporting durable objective response in 15% of patients, with 21% of patients remained progression-free at 1 year (22,23). Unfortunately, in March 2026 TZM was withdrawn from all global markets by the manufacturer following results from the Phase Ib/III SYMPHONY-1 trial in follicular lymphomas, which identified a potential risk of secondary hematologic malignancies. Despite this decision, EZH2 inhibition showed antitumor activity in patients with advanced EpS (22,23). A significant number of new EZH2i are being tested in clinical trials alone or in combination with other anti-cancer agents on different solid tumors (24–29). Additionally, innovative approaches, including the development of EZH1/EZH2 dual inhibitors and PROTAC degraders, are currently being investigated to target EZH2 in cancer (30,31). These considerations leave open the opportunity of future options to target EZH2 in EpS and foster investigations into mechanisms underpinning resistance to identify modifiers of tumor response and new treatment strategies, including treatment combinations.

Preclinical evidence has suggested that EZH2 blockade renders chromatin more accessible, thereby potentiating chemotherapy-induced DNA damage and apoptosis (32). In acute myeloid leukemia models, co-treatment with EZH2i and doxorubicin (DOX) induced transcriptional reprogramming and enhanced tumor suppression (33). Notably, the clinical rationale for combining EZH2i with DOX was pursued prospectively in a phase 1b/3 trial (ClinicalTrials.gov Identifier: NCT04204941) evaluating TZM in combination with DOX a frontline therapy for advanced EpS. However, the study was terminated early, before the randomized phase could be initiated, leaving the clinical activity of this combination formally unevaluated (34). Here, we modeled this therapeutic strategy in two in-house generated patient-derived xenografts (PDXs), and corresponding cell lines, of P-EpS, capturing two clinically relevant patterns of inadequate response – adaptive persistence and primary refractoriness. We showed that target engagement by EZH2 inhibition is not sufficient for efficacy, and that productive chromatin remodeling, marked by H3K27ac, distinguishes responsive from resistant tumor models. Pan-histone deacetylase (HDAC) inhibition partially restores this chromatin transition and sensitivity to the EZH2i–DOX combination, highlighting an actionable vulnerability with translational potential in INI1-deficient tumors.

## Materials and Methods

### Patient and human tumor characterization

Two INI1-deficient P-EpS samples suitable for mouse implantation were obtained from the primary tumor of two patients with naive primary EpS who underwent surgery.

The first model, named EpS-1, was previously described in (35). Briefly, this was a *SMARCB1*-deleted P-EpS arising in the forearm of a male patient, who developed disease progression after surgery. The second one, named EpS-2, was obtained from a 43-year-old male patient, with a diagnosis of *SMARCB1*-deleted P-EpS arising from the perineal region with concomitant bone metastasis. Immunohistochemistry (IHC) analysis showed loss of INI1 expression. This patient developed metastatic disease and eventually died within 8 months.

The use of patient material for PDX generation was approved by the institutional ethical committee (INT code: 16/17), and informed consent was obtained from all patients. Generation of PDX models and *in vivo* drug experiments were approved by the Institutional Animal Care and Use Committee and authorized by the Italian Ministry of Health (project approval code: 234/2018-PR), in accordance with international policies and national regulations (Directive 2010/63/EU).

### Development of P-EpS PDX and PDX-derived cell line

The EpS-1 and EpS-2 PDX models were generated from fresh P-EpS specimens collected immediately after surgical resection from these previously described patients, as reported elsewhere (35) and detailed in Supplementary Material S1. A third PDX model, designated EpS-1/R, was derived from EpS-1 PDX that resumed growth following TZM+DOX *in vivo* treatment cessation (see *Results* – *EZH2i–DOX combination shows robust antitumor activity in the EpS-1 PDX model, even after rechallenge)*.

EpS-1 and EpS-2 cell lines were generated from the corresponding PDXs, essentially as previously described (35).

Authentication of PDX models and corresponding cell lines was performed by microsatellite analysis using the AmpFISTR Identifier PCR Amplification Kit (Applied Biosystems; RRID: SCR_005039).

### Characterization of PDX models and cell lines

#### IHC analysis

Four-micrometer sections of formalin-fixed, paraffin-embedded (FFPE) tumor tissues from patients’ surgical specimens and xenografts were stained with hematoxylin and eosin (H&E) for morphological assessment, and immunostained for INI1 and Ki67. IHC was performed at room temperature using the Dako Autostainer Link 48 AS480 (Agilent; RRID: SCR_013575). Primary antibodies included anti-Ki67 (1:400; clone MIB-1, Agilent, Cat. #GA626; RRID: AB_2687921) and anti-INI1 (1:200; clone 25, Becton Dickinson, Cat. #612110; RRID: AB_399481).

Slides were processed with sequential incubations in EnVision Peroxidase (5 min, Dako), primary antibody (30 min), EnVision FLEX Linker (15 min), EnVision FLEX horseradish peroxidase (20 min), and diaminobenzidine (DAB; 10 min) for chromogenic detection, with buffer rinses between steps. Sections were counterstained with hematoxylin (2 min, Dako), rinsed, and coverslipped.

Digital images were analyzed using QuPath v0.1.2 (RRID: SCR_018257). The Ki67 labeling index was calculated as the percentage of Ki67-positive nuclei relative to total nuclei: (Ki67⁺ nuclei / total nuclei) × 100.

Sarcoma pathologists compared the morphological features of human tumors and corresponding PDX samples based on H&E and IHC analyses.

#### FISH analysis

*SMARCB1* gene status was evaluated using two dual-color FISH probes. The commercially available SPEC SMARCB1/22q12 Dual Color Probe (ZytoLight/ZytoVision, Bremerhaven, Germany) consisted of a 545-kb Spectrum Green-labeled probe targeting the *SMARCB1* gene (22q11.23) and a Spectrum Orange-labeled control probe targeting the *KREMEN1* gene (22q12.2). The in-house dual-color probe consisted of a 147-kb *SMARCB1*-specific Spectrum Green-labeled BAC probe (RP11-71G19) and a Spectrum Orange-labeled control probe (RP11-506F7) targeting the *PDGFB* locus (22q13.1) (35).

#### Genomic profile

Genomic DNA was isolated from FFPE sections of EpS PDXs using the GeneRead DNA FFPE Kit (Qiagen; RRID: SCR_008539), following the manufacturer’s recommendations. DNA concentration was quantified with the Qubit dsDNA High-Sensitivity Assay Kit (Thermo Fisher Scientific Inc.; RRID: SCR_008452), and DNA integrity was evaluated using the TapeStation 4200 system (Agilent; RRID: SCR_018435).

The OncoScan™ CNV Plus Assay (Thermo Fischer Scientific Inc.) was used to detect genome-wide copy number changes, including amplifications, deletions, and copy-neutral loss of heterozygosity (LOH), together with selected somatic variants, according to standard procedures and GRCh38/hg38 as reference genome. CNV data visualization and interpretation were performed using the karyoploteR R package (36).

#### DNA methylation profile

Genomic DNA was extracted by using the DNeasy Blood and Tissue Kit (Qiagen) following the manufacturer’s instructions. DNA (500ng) was bisulfite converted with the EZ DNA Methylation™ Kit (Zymo Research) and hybridized on Infinium MethylationEPIC v2.0 BeadChips (Illumina) (37). A NextSeq550 instrument (Illumina) was used for array scanning. Data analysis was conducted as previously described (13). t-SNE analysis was performed on beta values of the 10,000 most variable methylation probes using the Rtsne v 0.15 package. t-SNE including different data sets was performed on common probes, normalized, and batch-corrected as previously described (38). The additional data sets considered in the analysis were from our previously published EpS cohort and other studies, including clinical cases of EpS from the EPISObs observational study [Italian Sarcoma Group. *An observational study on epithelioid sarcoma (EPISObs)*. ClinicalTrials.gov Identifier: NCT03099681] and other STSs (DKFZ cohort) (13,38,39).

#### Transcriptomic profile

Total RNA was isolated from frozen PDX samples (3 biological replicates per PDX model) with a RNeasy mini kit (Qiagen). RNA quality was subsequently verified by measuring concentration and purity on a spectrophotometer, and determining the RNA integrity number (RIN) on an Agilent TapeStation 4200 (RNA ScreenTape Assay kit, RRID: SCR_018435, Agilent Technologies). Whole RNA-sequencing libraries were prepared from 250 ng total RNA using the Illumina TruSeq Stranded total RNA kit (Illumina) following manufacturer’s instructions, and sequenced on an Illumina HiSeq apparatus. The quality of the row sequencing data was evaluated with the FastQC software. Reads were preprocessed with fastp and aligned to both the human (GRCh38) and mouse (GRCm39) reference genomes by using STAR (40) with default parameters. To discriminate human and mouse sequences in PDX samples, disambiguate algorithm (41) was used to retain human sequences only. RSEM (42) was used to quantify reference transcript abundance.

#### Transcriptomic analysis

For RNA-sequencing datasets collected on PDX models at either baseline and post-treatment, raw counts with low expression were filtered using filterByExpr function, and genes lacking an official gene symbol were excluded. Reads mapping to the same gene symbol were collapsed by summing their counts. Data was normalized using the trimmed mean of M-values (TMM) method implemented into the edgeR package (43) within the R environment. For post-treatment samples only, also unwanted technical variation was additionally evaluated using RUVs function of the RUVSeq package (44). In this way, batch effect was estimated and removed for downstream analyses.

Additional public available samples were derived from our previously published EpS cohort (13), from The Cancer Genome Atlas (TCGA) sarcoma cohort (45), and from the dbGaP study (accession phs003188.v1.p1), including samples from patients treated at the Memorial Sloan Kettering Cancer Center (MSKCC, New York) collected within the EZH-202 phase II clinical trial (NCT02601950).

Focusing on protein-coding genes, t-SNE analysis was conducted using the Rtsne v0.15 package on the top 2,000 most variable protein-coding genes, based on log2-transformed pTPM values (log2[pTPM + 0.1]). Batch effects were corrected using the removeBatchEffect function from the limma package prior to dimensionality reduction (37).

For differential expression analysis, data heteroscedasticity was removed using the voom method from the edgeR package (43) and a linear model was applied using the limma package (46) with empirical Bayes moderation to identify transcriptomic changes, measured as both log_2_-transformed fold change (FC) and t-statistic. To consider multiple test comparisons, the false discovery rate (FDR) correction was applied, and 0.05 value was used to determine statistical significance threshold.

Over-representation analysis was performed by using the gProfiler package (47) on the main databases of pathways and annotation (Gene Ontology, KEGG, and Reactome). As input, significantly (FDR < 0.05) either down-regulated (log_2_(FC) < −2), or up-regulated (log_2_(FC) > 2) gene lists were chosen, while all genes of the array were selected as background. Default adjusted p-value correction was selected, and a threshold of 0.05 was chosen to assess significance.

Gene set enrichment analysis (GSEA) was performed on the different collection of the Molecular Signatures Database (MSigDB). Genes were ranked by their t-statistic, and the fgsea package (48) was used to assess significant enrichments with an FDR threshold of 0.1.

For single-sample GSEA (ssGSEA), a custom collection of gene sets was used to calculate the enrichment score in each sample using the GSVA R package in R, applying the ssGSEA method with normalization enabled.

### Growth inhibition assay

For *in vitro* studies, DOX (Adriblastina, Pfizer Inc., RRID: SCR_021375) was diluted in sterile saline solution, while TZM (EPZ-6438, MedKoo Biosciences), bafilomycin A1 (BafA1, Sigma-Aldrich, RRID: SCR_008988) and vorinostat (SAHA, Selleckchem, RRID: SCR_003823) were dissolved in DMSO and diluted in culture medium (0.5% final concentration).

For TZM–DOX growth inhibition assays, EpS-1 and EpS-2 cells were seeded at densities of 13,000 cells/cm² and 8,000 cells/cm², respectively, and treated 24 hours later with increasing drug concentrations for 144 hours. TZM was tested at 10µM, 30µM, 60µM, and 100µM, while DOX was used at 0.01µM, 0.05µM, 0.1µM and 0.5µM. Antiproliferative activity was assessed by cell counting using a Coulter Counter (Z Series Coulter Counter, Beckman Coulter GmbH, RRID: SCR_008940), and concentrations able to inhibit cell proliferation by 50% (IC_50_) were calculated from dose-response curves. For combination treatment, TZM IC_50_ concentrations (47µM and 38µM of TZM for EpS-1 and EpS-2 cells, respectively) were combined with 0.01µM, 0.05µM, 0.1µM and 0.5µM of DOX. Vehicle-treated cells (saline solution and/or DMSO 0.5%) were used as controls. Drug interactions were evaluated using the combination index (CI), where CI < 0.9, CI = 0.9–1.1, and CI > 1.1 indicated synergistic, additive, and antagonistic effects, respectively, using CalcuSyn software (Biosoft, RRID: SCR_020251).

For autophagy inhibition experiments, BafA1 (15nM) was combined with either TZM, DOX or their combination under the conditions described above, and cells were treated for 72 hours.

For triple-combination experiments, a three-dimensional dose matrix design was used, treating EpS-1 and EpS-2 cells for 72 hours with IC_50_, IC_30_, and IC_10_ concentrations of TZM (EpS-1: 50 µM, 31 µM, and 15 µM; EpS-2: 64 µM, 30 µM, and 13 µM), DOX (EpS-1: 0.09 µM, 0.03 µM, and 0.01 µM; EpS-2: 0.1 µM, 0.04 µM, and 0.01 µM), and SAHA (EpS-1: 8 µM, 4 µM, and 1 µM; EpS-2: 6 µM, 2.8 µM, and 1 µM), as either single agents or combination. Antiproliferative activity was assessed by cell counting, and drug interactions were quantified using the zero-interaction potency (ZIP) score calculated with the online tool SynergyFinder (RRID: SCR_026127) (49). For visualization of triple-drug interactions, SynergyFinder applies a multidimensional scaling-based dimensionality reduction approach that projects multidimensional dose combinations into a two-dimensional synergy landscape while preserving similarity among neighboring dose conditions (50).

For *in vivo* drug activity studies, tumor-bearing mice were randomized to treatment groups once PDX tumors reached approximately 150 mm³. Eight mice were used for each experimental group for EpS-1 model; six for EpS-1/R, and five for EpS-2 based on power calculation and growth rate. Group sizes were determined based on the experimental design and model-specific characteristics, including tumor growth kinetics and material availability. CD-1 NUDE (Crl:CD1-*Foxn1^nu^*, Charles River Laboratories, RRID: IMSR_CRL:086) mice were used for EpS-1 and EpS-1/R PDX models, whereas SCID mice were employed for EpS-2 due to lack of tumor engraftment in NUDE mice. TZM was formulated in 0.5% carboxymethylcellulose and 0.1% Tween 80 and administered orally (p.o.) at 250 mg/kg in EpS-1 and EpS-1/R models, while at 125 mg/kg in EpS-2, twice daily for 21 consecutive days (2qd x 21). DOX was dissolved in saline solution and administered intravenously (i.v.) every 7 days for three doses (q7d x 3) at 4 mg/kg in EpS-1 and EpS-1/R models, while 2.5 mg/kg in EpS-2. The same schedules were applied for single-agent and combination treatments. Vehicle-treated mice served as controls. Drug activity was evaluated as TV inhibition (TVI) percentage using the formula

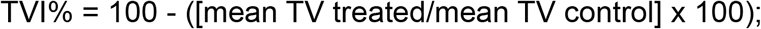

Treatment tolerability was determined by monitoring body weight loss and lethal toxicity. For transcriptomic analysis of treated PDX tumors, three SCID mice were used for each experimental group for both EpS-1 and EpS-2 PDX models. TZM was administered orally (p.o.) at 250 mg/kg twice daily for 8 consecutive days (2qd × 8), while DOX was given intravenously (i.v.) at 2.5 mg/kg every 7 days for two doses (q7d × 2). This short-term *in vivo* treatment schedule was designed to capture early transcriptional changes associated with on-target drug activity.

### Western Blot analysis

Lysates were obtained from frozen PDX tumors collected immediately after the end of treatment and from the PDX-derived cell lines at different intervals after exposure to each drug. PDX tumor tissues were pulverized using the Mikro-Dismembrator II (B. Braun Biotech International) prior to lysis. All samples were lysed in 125 mM Tris-HCl (pH 6.8), 5% SDS, and 25 mM NaF, supplemented with 1X Halt Protease and Phosphatase Inhibitor Cocktail (Thermo Fischer Scientific, #78443) and protein concentration was quantified using the Pierce™ BCA Protein Assay Kit (Thermo Fisher Scientific, #A65453). Equal amounts of proteins were separated by SDS-PAGE, transferred onto nitrocellulose membranes, and incubated with the following primary antibodies: anti-ß-actin, anti-ß-catenin, anti-cleaved-caspase-3, anti-E-cadherin, anti-EZH2, anti-GAPDH, anti-histone H3, anti-H3K27ac, anti-H3K27me^3^, anti-LC3B, anti-PARP, anti-INI1, anti-ß-tubulin, anti-vimentin, anti-vinculin. To detect primary antibodies, horseradish peroxidase-linked goat anti-rabbit IgG or horseradish peroxidase-linked horse anti-mouse IgG antibodies were employed. Antibody details are reported in Supplementary Table S1. The enhanced chemiluminescence detection system (ECL, GE Healthcare, RRID: SCR_000004) was used. To allow the simultaneous detection of multiple proteins with different molecular weights, the membranes were cut into different parts. Membranes were stripped and re-incubated with different primary antibodies, followed by incubation with the appropriate horseradish peroxidase–conjugated secondary antibodies. For figure preparation, original Western blot films were cropped and assembled to generate composite panels showing different proteins. Molecular weight markers were visualized using the Precision Plus Protein Standard (Bio-Rad Laboratories, RRID: SCR_008426). The quantification of all the band’s intensities is reported in Supplementary Tables.

### Apoptosis analysis

Cell apoptosis was evaluated by Terminal deoxynucleotidyl transferase dUTP Nick-End Labeling (TUNEL) assay by using the In Situ Cell Death Detection Kit (Sigma-Aldrich, RRID: SCR_008988) according to manufacturer’s protocol. Briefly, EpS-1 and EpS-2 cells were seeded at densities of 13,000 cells/cm² and 8,000 cells/cm², respectively, in 25 cm^2^ flasks and treated 24 hours later with TZM and DOX at their IC_50_ concentrations, either as single agents or in combination, for 72 or 144 hours. For autophagy inhibition experiments, BafA1 was used at 15nM, and cells were treated for 72 hours. Vehicle-treated cells (saline solution and/or DMSO 0.5%) served as controls. At the end of treatment, both floating and adherent cells were collected for apoptosis analysis. Cells were fixed in freshly prepared 4% paraformaldehyde for 45 min at room temperature and permeabilized with 0.1% Triton X-100 in 0.1% sodium citrate for 3 min on ice. DNA strand breaks were labeled by incubation with TUNEL reaction mixture for 1 hour at 37°C in a humified atmosphere and protected from light. After washing, samples were analyzed by flow cytometry using BD Accuri C6 Plus flow citometer (BD Biosciences, RRID: SCR_019591) and data were processed with Kaluza analysis software (Beckman Coulter, RRID: SCR_016182).

A detailed description for caspase-3 catalytic activity assay is reported in Supplementary Material S1.

### Migration and invasion

Cell migratory capacity was evaluated using a wound healing assay by measuring the ability of cells to close a gap in the culture monolayer. EpS cell lines were seeded into two-well silicone culture inserts with a defined cell-free gap (ibidi, 80209). Twenty-four hours after seeding, inserts were removed, and cells were supplied with fresh complete medium. Phase-contrast images were acquired immediately after insert removal (*t_0_*) and after 24 (*t_24_* h) and 48 hours (*t_48_* h). After imaging the wound at the indicated time points, the wound area was measured using ImageJ software (version 1.47q; RRID: SCR_003070). Migration ability was expressed as the percentage of wound closure.

Cell invasive capacity was evaluated using Transwell permeable supports (Corning, #3422, 8.0 μm polycarbonate membrane, 6.5 mm inserts) pre-coated with Matrigel (Corning, #356231). EpS cell lines were plated in the upper chamber in serum-free medium, while the lower chamber was supplemented with complete medium. Twenty-four hours later, cells that had invaded the Matrigel and migrated across the porous membrane were fixed with 100% ethanol, stained with 0.4% sulforhodamine B (Merck) in 1% acetic acid, and subsequently washed. Images from five random fields per chamber were acquired using phase-contrast microscopy and quantitatively analyzed with ImageJ.

### Statistical analysis

Statistical analysis was performed with Mann-Whitney test and Kruskal–Wallis test (Dunn’s method for multiple comparisons), when appropriate, using GraphPad Prism software (version 9.4; GraphPad Prism Inc., San Diego, CA, USA). The TumGrowth model (https://kroemerlab.shinyapps.io/TumGrowth) was employed to examine the effect of treatment and time on cell viability and TVI, including random effects to account for interanimal variability. The threshold for statistical significance was set to 0.05.

### Data availability

Baseline transcriptomic data from EpS-1 and EpS-2 PDX were deposited together with transcriptomic profiles of EpS-1 and EpS-2 PDX after treatment with either DOX, TZM or their combination at Gene Expression Omnibus (accession number GSE339069 and GSE327877, respectively). All other relevant data generated during the current study are available from the corresponding author on reasonable request.

## Results

### Patient-derived model generation and characterization

A P-EpS PDX named EpS-1 has been established by our group and utilized in a previous study (35). An additional model, EpS-2, was generated from another patient with P-EpS. The newly generated PDX recapitulated the histomorphology of the patient’s clinical tumor, as confirmed by H&E staining where the tumor exhibits a nodular growth pattern with cystic and hemorrhagic components. Both clinical tumor and PDX showed variable cellularity, ranging from low- to high-density areas, display myxoid stroma, necrosis, smaller tumor cell nuclei compared to EpS-1, and rhabdoid features (**Fig. 1A**). A complete loss of INI1 nuclear immunohistochemical staining was observed, consistent with the homozygous deletion of the *SMARCB1* gene detected by FISH analysis in both the patient tumor and the EpS-2 PDX (**Fig. 1A**). EpS-2 cells, which were derived from the paired PDX model, exhibited larger size, increased cytoplasmic volume, and a spindle-shaped morphology compared with the more cuboidal appearance of EpS-1 cells (**Fig. 1B**). Consistently with the parental PDX tumors, FISH analysis confirmed a homozygous deletion of the *SMARCB1* gene in both EpS-1 and EpS-2 cell lines (**Fig. 1B**). Absence of INI1 was also confirmed by Western blot analysis in both PDXs and cell lines (Supplementary Fig. S1C).

**Figure 1.**
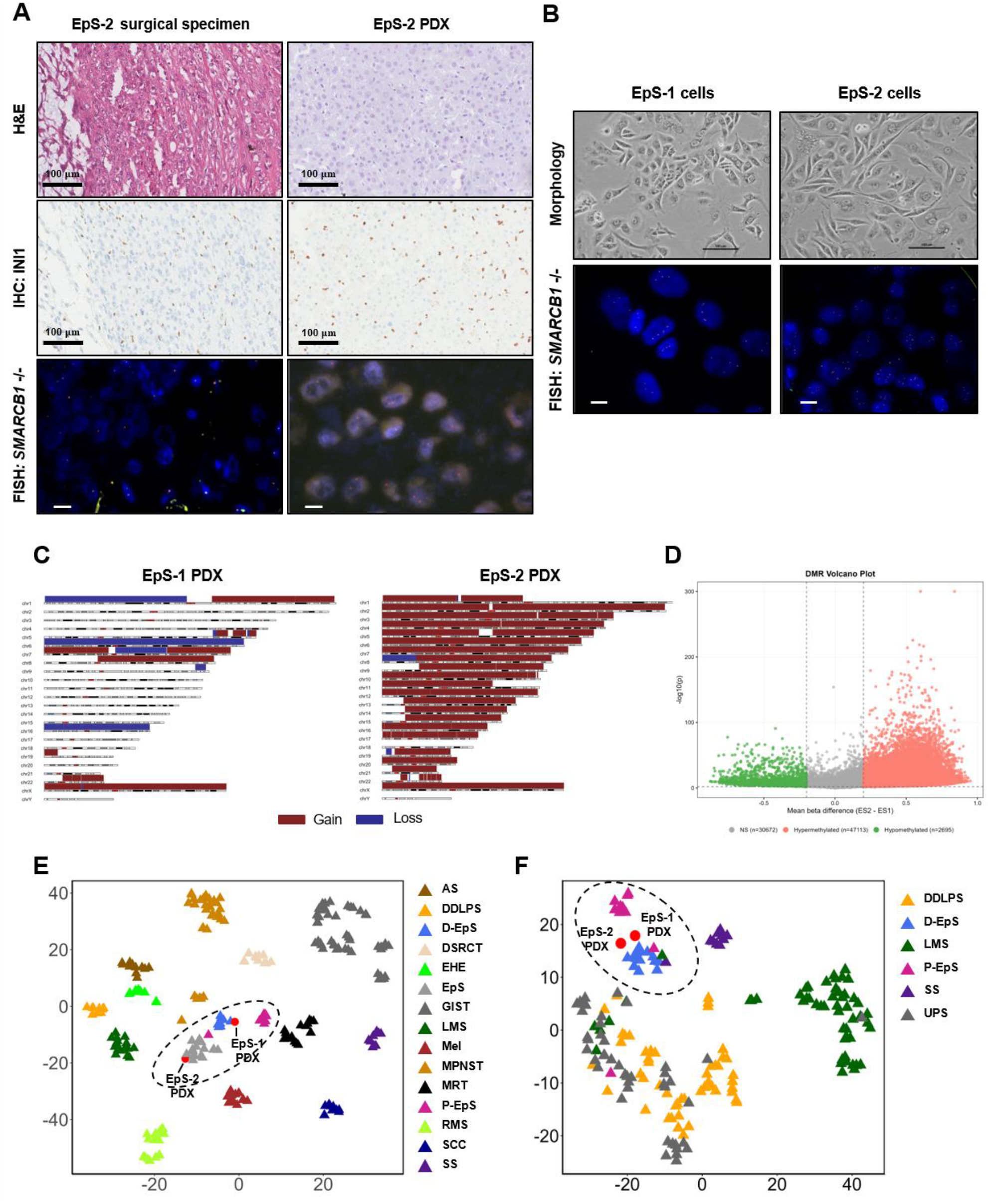
Characterization of EpS patient-derived xenograft (PDX) models and paired cell lines. **A**, Histological features were evaluated on haematoxylin and eosin (H&E)-stained sections (upper panels; scale bars, 100 µm). INI1 protein expression was assessed by immunohistochemistry (IHC) (middle panels; scale bars, 100 µm). FISH analysis using the ZytoLight/ZytoVision SPEC SMARCB1/22q12 Dual Color Probe showed *SMARCB1* homozygous deletion in both the EpS-2 clinical specimen and the corresponding PDX model. *SMARCB1*-retaining non-neoplastic cells served as internal controls in the surgical specimen (lower panels; scale bars, 10 µm). **B**, Morphology of EpS-1 and EpS-2 cell lines derived from the corresponding PDX models, growing as monolayers (upper panels; scale bars, 100 µm). FISH analysis using the in-house dual-color probe (SMARCB1/PDGFB) showed complete loss of *SMARCB1* signals in neoplastic cells (lower panels; scale bars, 10 µm). **C**, Genomic profiles of EpS-1 and EpS-2 PDX models, showing copy number losses (blue) and gains (red). *SMARCB1* loss is present in both models. **D**, Volcano plot of differentially methylated DNA regions at baseline (EpS-2 PDX vs EpS-1 PDX). **E**, Scatterplot of t-distributed stochastic neighbor embedding (t-SNE) analysis of DNA methylation profiles from the metadataset samples. **F**, Scatterplot of t-SNE analysis of transcriptomic profiles from the metadataset samples. AS, angiosarcoma; DDLPS, dedifferentiated liposarcoma; D-EpS, distal epithelioid sarcoma; DSRCT, desmoplastic small-round cell tumor; EHE, epithelioid hemangioendothelioma; EpS, epithelioid sarcoma; GIST, gastrointestinal stromal tumor; LMS, leiomyosarcoma; Mel, melanocytes; MPNST, malignant peripheral nerve sheath tumor; MRT, malignant rhabdoid tumor; P-EpS, proximal epithelioid sarcoma; RMS, rhabdomyosarcoma; SCC, squamous cell carcinoma; SS, synovial sarcoma; UPS, undifferentiated pleomorphic sarcoma.

Genome-wide copy-number analysis highlighted marked differences in the chromosomal architecture of the two PDX models (**Fig. 1C**, Supplementary Fig. S2). While EpS-1 displayed a more focal pattern of copy-number alterations, the EpS-2 PDX was characterized by genomic imbalance, dominated by broad arm-level and chromosome-wide gains affecting multiple autosomes. Deletion of the *SMARCB1* locus was confirmed in both models. Collectively, these features define EpS-2 as a PDX model with widespread chromosomal gains and increased copy-number complexity compared with EpS-1.

To assess whether the PDX models faithfully recapitulated the transcriptional and epigenetic landscape of the original histotype, t-SNE analysis based on genome-wide DNA methylation and transcriptomic profiles was conducted. The analyses showed that, although EpS-2 DNA was hypermethylated compared with EpS-1 DNA (**Fig. 1D**), both PDX models clustered with clinical EpS (**Fig. 1E**). Similarly, t-SNE of the transcriptomic data showed that EpS-1 and EpS-2 spanned the space between tumors from patients who were classified as having a D- or a P-EpS (**Fig. 1F**). These data suggest that, despite their molecular differences, the PDX models faithfully recapitulate the overall epigenetic and transcriptional features of clinical EpS.

### EZH2i plus DOX shows distinct interaction patterns between EpS cell lines

To assess the cytotoxic effects of TZM, DOX, and their combination, EpS-1 and EpS-2 cell lines were treated with each agent alone or in combination for 144 hours. Both TZM and DOX induced dose-dependent growth inhibition, with DOX displaying greater activity than TZM, as indicated by the lower concentrations required to achieve 50% growth inhibition (IC_₅₀_: 0.04 µM vs. 46 µM in EpS-1 cells; 0.05 µM vs. 39 µM in EpS-2 cells, respectively). Combined treatment with TZM and DOX resulted in enhanced growth inhibition compared with either single agent alone (**Fig. 2A**). Notably, when TZM was administered at its IC_₅₀_ concentration in combination with increasing concentrations of DOX, the interaction between the two agents differed between the two cell lines. Specifically, the combination exerted synergistic effects in EpS-1 cells (CI < 1), whereas an antagonistic interaction was observed in EpS-2 cells (CI > 1) (**Fig. 2B**).

**Figure 2.**
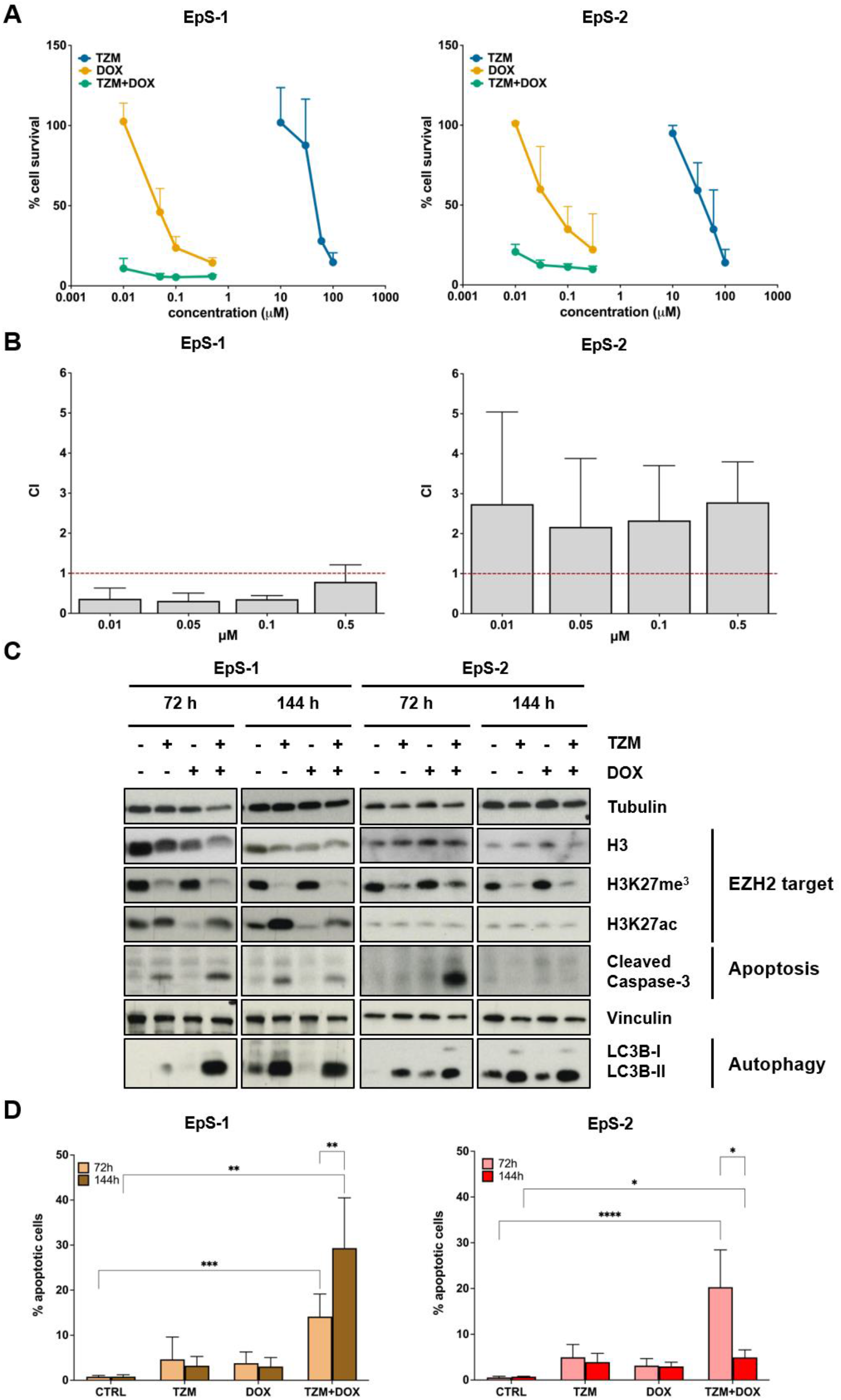
Tazemetostat (TZM)-doxorubicin (DOX) combination exhibited distinct interaction profiles in EpS-1 and EpS-2 cell lines. **A**, Cell growth inhibition curves following treatment with TZM, DOX, or their combination in EpS-1 and EpS-2 cells. Data are expressed as percentage of viable cells relative to untreated controls and represent mean ± SD of three independent experiments. **B**, Combination index (CI) plots describing the interaction between TZM and DOX in EpS-1 and EpS-2 cells. Data represent mean ± SD of three independent experiments. **C**, Western blot analysis of single agent and combined treatment-induced biological effects on EZH2 downstream targets, apoptosis, and autophagy in EpS-1 and EpS-2 cells. Representative cropped images are shown. Tubulin and vinculin were used as loading controls. **D**, Quantification of TUNEL-positive cells assessed by flow cytometry after treatment with single agents or the combination in EpS-1 and EpS-2 cell lines. Data represent mean ± SD of three independent experiments. Statistical analysis: comparisons among treatment groups at each time point were performed using the Kruskal-Wallis test with Dunn’s multiple comparison test. The comparison between 72 and 144 hours within the TZM+DOX group was performed using the Mann-Whitney test. *p < 0.05; **p < 0.01; ***p < 0.001; ****p < 0.0001.

To investigate the biological effects of single-agent and combination treatments, Western blot analyses were performed after 72 and 144 hours of drug exposure in both EpS-1 and EpS-2 cell lines (**Fig. 2C**, Supplementary Table S2). To verify the on-target activity of TZM, we first assessed the levels of H3K27 trimethylation (H3K27me^3^). Treatment with TZM, either alone or in combination with DOX, effectively reduced H3K27me^3^ levels in both cell lines. Given the dynamic interplay between repressive and activating histone modifications at H3K27, we additionally evaluated H3K27 acetylation (H3K27ac) following EZH2 inhibition. Notably, TZM, alone or in combination with DOX, induced an increase in H3K27ac levels exclusively in EpS-1 cells. Interestingly, TZM treatment, with or without DOX, induced apoptosis at both time points in EpS-1 cells, as evidenced by caspase-3 cleavage, whereas in EpS-2 cells the combination treatment triggered apoptosis only after early drug exposure (**Fig. 2C**, Supplementary Table S2). Differential apoptosis induction following combined treatment in the two cell lines was further supported by flow cytometry analysis, which revealed that while an increased proportion of TUNEL-positive cells was observed in both EpS-1 and EpS-2 cell lines at 72 hours, this effect was maintained only in EpS-1 cells at 144 hours, in agreement with the Western blot results (**Fig. 2D**). To further substantiate the induction of caspase-dependent apoptosis, caspase-3 catalytic activity was measured in EpS cell lines following single-agent and combined treatments. Treatment with either TZM or DOX alone resulted in minimal changes in caspase-3 activity in both cell lines. In contrast, combined TZM and DOX treatment led to an increase in caspase-3 activity at 72 hours in both EpS-1 and EpS-2 cells, which was maintained at 144 hours in EpS-1 cells only, consistent with the Western blot and TUNEL results (Supplementary Fig. S3). Based on the established role of EZH2 in the epigenetic regulation of autophagy (51), and in light of previous evidence showing autophagy induction following EZH2 inhibition with the TZM analogue EPZ-011989 (35), we assessed autophagy-associated changes by analyzing the kinetics of LC3B conversion after drug treatment. Accumulation of LC3B-II was observed following TZM treatment, either alone or in combination with DOX, in both cell lines at both time points, indicating a role of autophagy in TZM response as well (**Fig. 2C**, Supplementary Table S2).

### TZM-induced autophagy has a cytoprotective role in the EpS-2 cell line

Based on the observation that TZM, either alone or in combination with DOX, induced an autophagic response in both EpS-1 and EpS-2 cells in spite of a different susceptibility to the combined treatment of the two cell lines, we next explored the functional contribution of autophagy to the cellular response to treatments. This analysis was further prompted by the distinct temporal patterns of apoptotic activation observed in the two models, characterized by a prolonged apoptotic response in EpS-1 cells and a transient apoptotic response in EpS-2 cells following combined treatment. To this end, EpS-1 and EpS-2 cells were treated with TZM and DOX, alone or in combination, in the presence or absence of the autophagy inhibitor BafA1, a macrolide antibiotic that blocks late-stage autophagy by preventing autophagosome–lysosome fusion (52), and cell growth inhibition was assessed after 72 hours. In EpS-1 cells, autophagy inhibition did not modify drug response, whereas BafA1 significantly increased the cytotoxic effects of TZM and its combination with DOX in EpS-2 cells (**Fig. 3A**).

**Figure 3.**
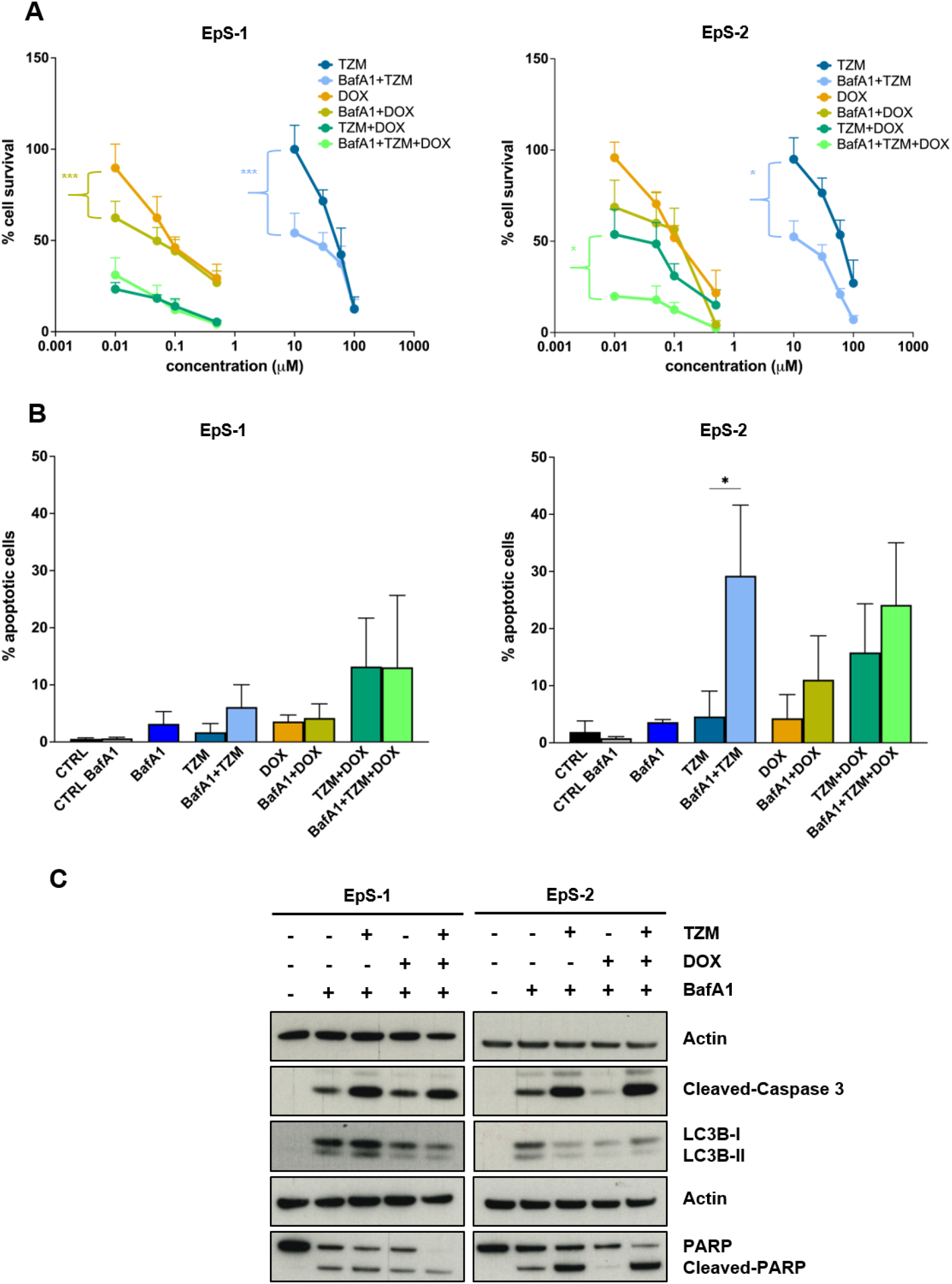
Tazemetostat (TZM)-induced autophagy exerts a cytoprotective effect in EpS-2 cells. **A,** Cell growth inhibition curves following treatment with TZM, doxorubicin (DOX), or their combination in the presence of Bafilomycin A1 (BafA1) in EpS-1 and EpS-2 cells. Data are expressed as percentage of viable cells relative to untreated controls and represent mean ± SD of three independent experiments. Statistical analysis: TumGrowth model. *p < 0.05, ***p < 0.001. **B,** Quantification of TUNEL-positive cells by flow cytometry after treatment with TZM, DOX, or their combination in the presence of BafA1 in EpS-1 and EpS-2 cell lines. Data represent mean ± SD of three independent experiments. Statistical analysis: Mann–Whitney test. *p < 0.05. **C,** Western blot analysis of apoptosis-related (cleaved caspase-3 and PARP) and autophagy-related markers (LC3B-I to LC3B-II conversion) in EpS-1 and EpS-2 cells treated with TZM and DOX, as single agents or in combination, with or without BafA1. Representative cropped images are shown. Actin was used as loading control.

To determine whether the increased growth inhibition observed upon autophagy blockade was associated with enhanced apoptotic activation, TUNEL assay was performed. Consistently, BafA1 significantly increased apoptosis following TZM-based treatments in EpS-2 cells, whereas no appreciable effects were detected in EpS-1 cells (**Fig. 3B**).

At the molecular level, BafA1 inhibition impaired LC3B processing and was associated with a marked increase in caspase-3 cleavage in EpS-2 cells following TZM (± DOX) exposure, reaching levels comparable to those observed in EpS-1 cells. Moreover, autophagy inhibition selectively induced PARP cleavage in EpS-2 cells after TZM and combined treatments, further supporting the engagement of apoptotic pathways upon autophagy blockade (**Fig. 3C**, Supplementary Table S3). Collectively, these findings indicate that autophagy contributes to limiting the cytotoxic effects of EZH2 inhibition in EpS-2 cells, consistent with a context-dependent, cytoprotective role of this process.

### EZH2i–DOX combination shows robust antitumor activity in the EpS-1 PDX model, even after rechallenge

To determine whether chromatin remodeling induced by EZH2i could enhance DOX cytotoxicity *in vivo*, EpS-1 and EpS-2 PDX models were treated with each agent alone or in combination. In the EpS-1 model, single-agent TZM and DOX produced modest antitumor effects, with maximum TVI values of 56% and 50%, respectively. In contrast, the combination treatment elicited pronounced antitumor activity, reaching a maximum TVI of 92% (**Fig. 4A and 4B**). The enhanced efficacy of TZM+DOX was supported by Ki67 immunostaining, which showed a reduction of the proliferation index to 26.6% compared with 81.9% in controls (**Fig. 4C**, Supplementary Fig. S4A). Morphological analysis of H&E sections corroborated these results: the untreated EpS-1 PDX displayed the typical epithelioid morphology with focal areas of basal necrosis, while combination-treated tumors showed increased myxoid stromal accumulation and more loosely connected neoplastic cells, suggestive of a pre-necrotic condition (Supplementary Fig. S4A).

**Figure 4.**
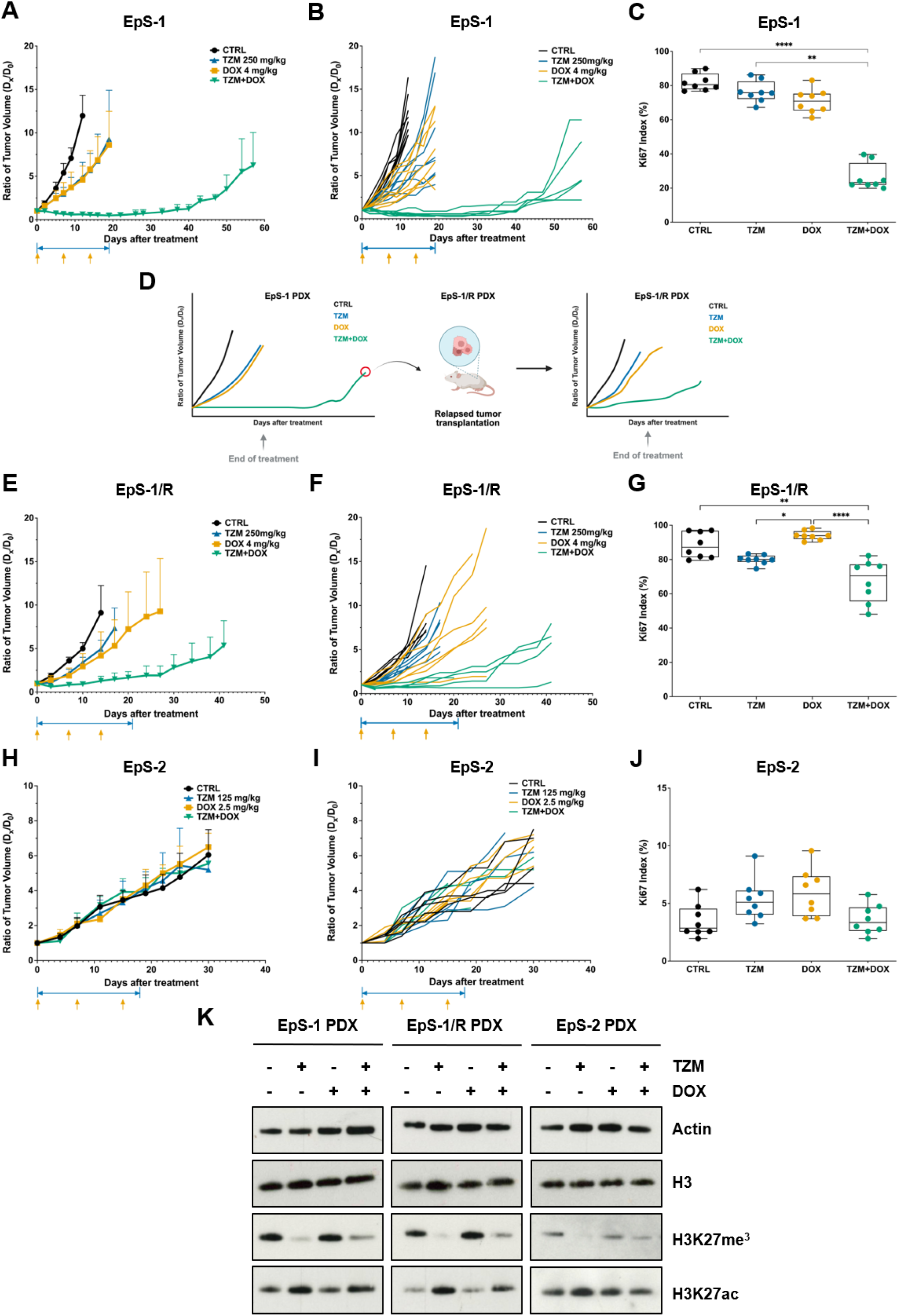
*In vivo* activity of tazemetostat (TZM), doxorubicin (DOX), and their combination. **A,** EpS-1 tumor growth curves expressed as the ratio of tumor volume (TV) at day x (D_x) relative to baseline (day 0, D_0) in control and TZM-, DOX-, and TZM+DOX-treated groups (n = 8). Arrows indicate drug administration. Data are mean ± SD. Statistical analysis: TumGrowth model. TZM vs CTR *, TZM+DOX vs CTR ****. *p < 0.05, ****p < 0.0001. **B,** Individual tumor growth curves, calculated from caliper measurements in EpS-1 patient-derived xenograft (PDX)-bearing mice treated with the indicated regimens. Arrows indicate drug administration. **C,** Quantification of Ki67 proliferation index in EpS-1 tumors collected at the end of treatment. Data represent mean ± SD from eight independent tumor areas. Statistical analysis: Kruskal–Wallis test. **p < 0.01, ****p < 0.0001. **D,** Schematic overview of the in vivo rechallenge experiment performed in the EpS-1/R PDX model derived from TZM-DOX-treated EpS-1 tumors after regrowth following treatment cessation. **E,** EpS-1/R tumor growth curves expressed as TV ratio relative to baseline in control and treated groups (n = 6). Data are mean ± SD. Statistical analysis: TumGrowth model. TZM+DOX vs CTR **. **p < 0.01. **F,** Individual tumor growth curves, calculated from caliper measurements in EpS-1/R PDX-bearing mice treated with the indicated regimens. Arrows indicate drug administration. **G,** Quantification of Ki67 index in EpS-1/R tumors at treatment end. Data represent mean ± SD from eight independent tumor areas. Statistical analysis: Kruskal–Wallis test. *p < 0.05, **p < 0.01, ****p < 0.0001. **H,** EpS-2 tumor growth curves expressed as TV ratio relative to baseline in control and treated groups (n = 5). Data are mean ± SD. **I,** Individual tumor growth curves, calculated from caliper measurements in EpS-2 PDX-bearing mice treated with the indicated regimens. Arrows indicate drug administration. **J,** Quantification of Ki67 index in EpS-2 tumors at treatment end. Data represent mean ± SD from eight independent tumor areas. **K,** Western blot analysis of histone H3, H3K27 trimethylation (H3K27me³) and H3K27 acetylation (H3K27ac) in tumors collected at the end of in vivo treatment. Cropped images of selected proteins are shown. Actin served as loading control.

EpS-1 tumors treated with TZM+DOX resumed growth approximately three weeks after treatment cessation. To evaluate the efficacy of a rechallenge therapy, a relapsed PDX model (EpS-1/R) was generated from regrowing EpS-1 tumors, along with its corresponding cell line (**Fig. 4D**). INI1 loss in EpS-1/R PDX and cell line was consistent with the parental EpS-1 model (Supplementary Fig. S1A–1C). In EpS-1/R, TZM showed slightly reduced activity (max TVI 40%), while DOX retained similar efficacy as in EpS-1 (max TVI 57%). Their combination remained more effective than single agents, achieving a max TVI of 84% (**Fig. 4E and 4F**), and Ki67 positivity was reduced to 67.5% versus 88.4% in controls (**Fig. 4G**, Supplementary Fig. S4B). Histologically, EpS-1/R mirrored the parental EpS-1 model, with combination treatment enhancing myxoid stromal areas and dispersed tumor cells, indicative of a pre-necrotic state not observed with single agents (Supplementary Fig. S4B). Unlike EpS-1, EpS-2 PDX was engrafted in SCID mice due to growth limitations in NUDE mice, and drug doses were reduced accordingly, to account for the greater susceptibility of SCID mice to treatment-related toxicity (53). In this model, neither single agents nor combination exerted inhibitory effects on tumor growth (**Fig. 4H and 4I**). Nonetheless, H3K27me^3^ was nearly completely inhibited, confirming effective EZH2 targeting by TZM, as observed in EpS-1 and EpS-1/R. However, consistent with the *in vitro* findings, no increase in H3K27ac was detected in the EpS-2 model following treatment, suggesting a distinct chromatin remodeling response to TZM-based therapies (**Fig. 4K**, Supplementary Table S4). Morphologically, EpS-2 tumors were predominantly composed of spindle-shaped cells, in contrast to the epithelioid morphology of EpS-1 and EpS-1/R, consistent with the slower growth kinetics of this PDX (**Fig. 4J**, Supplementary Fig. S4C).

### The resistant EpS-2 model exhibits an enriched epithelial-mesenchymal transition-like (EMT-like) gene expression program

To gain insight into the molecular mechanisms underlying the differences observed between the two models, RNA sequencing was performed on baseline PDX tumors. Transcriptomic profiles of EpS-1 and EpS-2 clustered distinctly in t-distributed stochastic neighbor embedding (t-SNE) space, whereas EpS-1/R samples localized close to the parental EpS-1 model (Supplementary Fig. S5A). Accordingly, EpS-1 and EpS-1/R PDXs displayed highly similar transcriptional profiles, with only a small number of differentially expressed genes (Supplementary Fig. S5B). In contrast, EpS-1 and EpS-2 PDXs showed extensive transcriptional divergence, with EpS-2 exhibiting 1429 significantly upregulated and 1649 significantly downregulated genes relative to EpS-1 (**Fig. 5A**). Overrepresentation analysis identified a marked enrichment of EMT-related processes in EpS-2, including cell motility, migration, extracellular matrix organization, and matrix stiffness (**Fig. 5B**, Supplementary Fig. S5C). Consistently, GSEA confirmed increased expression of EMT hallmark genes in the EpS-2 model (**Fig. 5C**).

**Figure 5.**
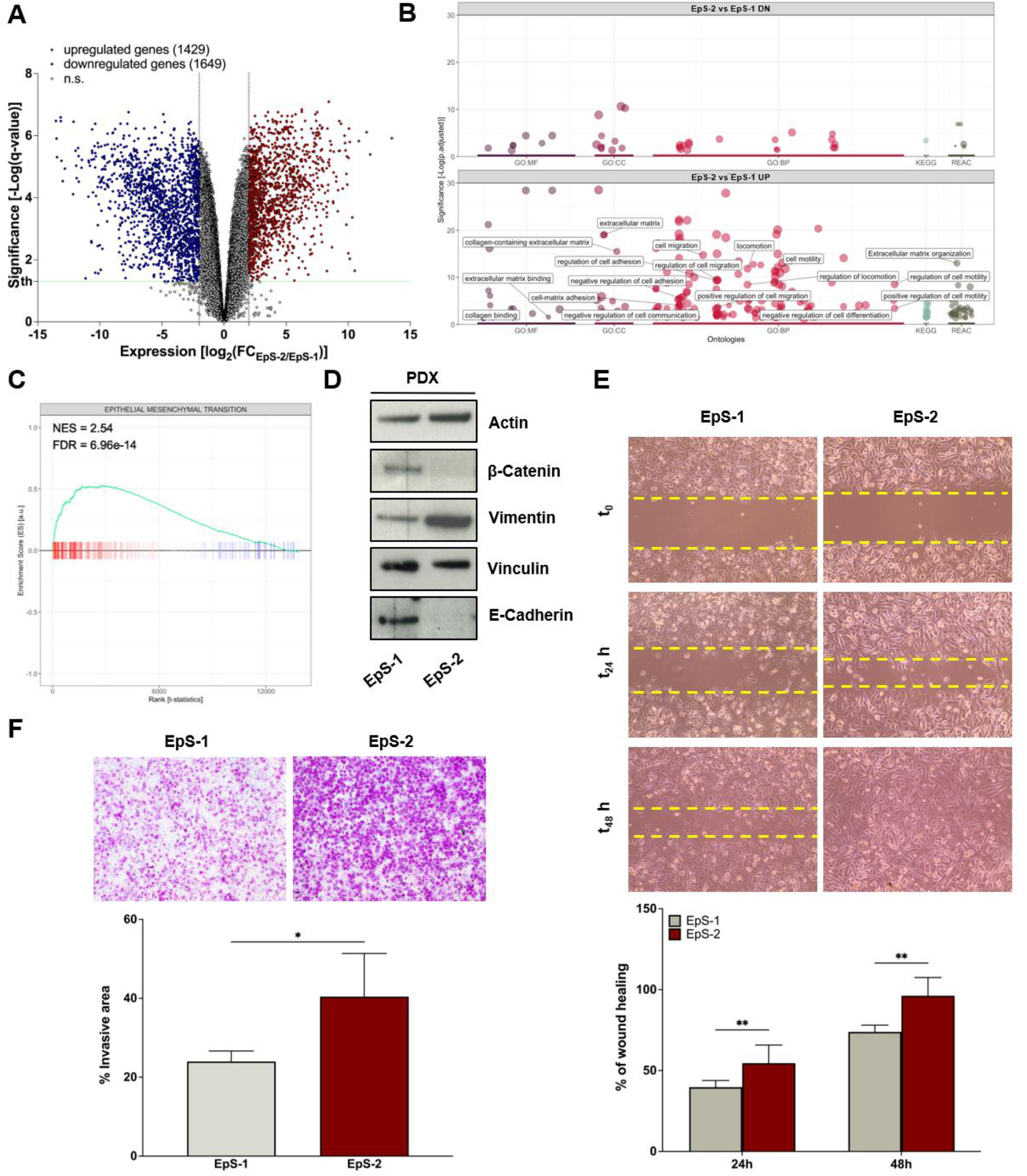
Baseline enrichment of epithelial-mesenchymal transition-like (EMT-like) transcriptional programs characterize the resistant EpS-2 PDX model. **A,** Volcano plot depicting differential gene expression between EpS-1 and EpS-2 PDX models at baseline. **B,** Manhattan plot showing significantly enriched gene sets identified by over-representation analysis (EpS-2 vs EpS-1 PDX models) across multiple functional databases. Each dot represents a gene set, grouped by database and plotted according to their enrichment significance. Named points indicate EMT-related biological processes. Gene Ontology (GO) with all the three ontologies: Molecular Function (MF), Cellular Component (CC), and Biological Process (BP), KEGG Pathway, and Reactome (REAC). **C,** Enrichment plot of Gene Set Enrichment Analysis (GSEA) of the Hallmark Epithelial–Mesenchymal Transition gene set in the EpS-2 PDX model (NES = 2.54; FDR = 6.96 × 10⁻¹⁴). **D,** Western blot analysis of epithelial markers (β-catenin and E-cadherin) and mesenchymal markers (vimentin) in EpS-1 and EpS-2 PDX tumors at baseline. Cropped images of selected proteins are shown. Actin and vinculin were used as loading controls. **E,** Representative bright-field micrographs illustrating the migratory capacity of EpS-1 and EpS-2 cells. Scale bar, 100 μm. Dotted lines indicate the cell-free wound area. Bar graphs (below) show wound closure percentages at 0, 24, and 48 hours. Data are mean ± SD. Statistical analysis: Mann–Whitney test. **p < 0.01. **F,** Representative bright-field micrographs showing the invasive capacity of EpS-1 and EpS-2 cells at 24 hours. Scale bar, 100 μm. Bar graphs (below) report the percentage of invaded area. Data are mean ± SD. Statistical analysis: Mann– Whitney test. *p < 0.05.

EMT is a reversible process whereby cells, including those of mesenchymal origin, such as sarcomas, can further enhance mesenchymal traits, becoming more motile, invasive, and resistant to radiation and chemotherapy (54–56). To validate these transcriptional findings, we assessed the expression of epithelial and mesenchymal markers in the two models. In line with RNA-seq data, EpS-1 PDXs displayed higher levels of β-catenin and E-cadherin, indicative of more epithelial-like features, whereas EpS-2 PDXs showed increased expression of vimentin, consistent with a more mesenchymal phenotype (**Fig. 5D**, Supplementary Table S5). Reflecting this differential phenotype, EpS-2 cells exhibited greater migratory ability, as indicated by the significantly increased wound closure, and invasive capacity compared with EpS-1 cells (**Fig. 5E and 5F**).

### HDAC inhibition restores H3K27ac and overcomes resistance to EZH2i-based therapy

To further investigate potential mechanisms underlying resistance to the TZM–DOX combination, RNA sequencing was performed on EpS-1 and EpS-2 tumors excised from mice after short-term *in vivo* treatment (Supplementary Fig. S6A). GSEA revealed that TZM-DOX modulated transcriptional programs in the EpS-1 PDX model, whereas no significant transcriptional reprogramming was detected in the resistant EpS-2 model (Supplementary Fig. S6B). In EpS-1, the TZM-DOX combination, but not either agent alone, resulted in a coordinated downregulation of multiple gene sets related to chromatin structure and epigenetic regulation, including HDAC-associated pathways, chromatin and chromosome organization, and MLL3/MLL4-dependent transcriptional control. Concomitantly, pathways involved in DNA metabolic processes and chromosome maintenance were also downregulated (**Fig. 6A**). Overall, these data suggest that sensitivity to TZM-DOX is associated with impairment of chromatin-dependent transcriptional and DNA regulatory programs. To explore whether this transcriptional signature had clinical correlates, we performed ssGSEA on publicly available transcriptomic data from a subset of EpS patients enrolled in the EZH-202 phase II trial of TZM (23,57). Among the six patients with available post-treatment transcriptomic profiles, the two TZM responders showed lower enrichment scores for chromatin organization-, HDAC-, and DNA metabolism-related gene sets compared with non-responding cases (**Fig. 6B**). Notably, both responders were primary tumors, whereas non-responders were derived from locally recurrent disease. Although the limited sample size and the heterogeneity of the clinical specimens preclude definitive conclusions, these findings support the hypothesis that effective EZH2 inhibition is associated with the disruption of chromatin-dependent transcriptional and DNA regulatory networks. The lack of increased H3K27ac following TZM-based treatments in the EpS-2 model, together with the transcriptional repression of chromatin remodeling and HDAC-related pathways observed in the responsive EpS-1 model and mirrored in a subset of TZM-responsive clinical samples, supported the hypothesis that HDAC-mediated chromatin regulation may contribute to resistance to EZH2 inhibition-based therapies. To functionally validate this hypothesis, EpS-1 and EpS-2 cells were treated with a triple combination of TZM, DOX, and the pan-HDAC inhibitor SAHA for 72 hours. The addition of SAHA enhanced the antiproliferative effects of TZM and DOX, both as single agents and in combination, in both cell lines (**Fig. 6C and 6D**). The triple combination resulted in a marked reduction of EZH2 protein levels, with a decrease in H3K27me^3^ and a concomitant increase in H3K27ac, ultimately leading to apoptotic cell death as indicated by cleavage of caspase-3 (**Fig. 6E and 6F**, Supplementary Table S6). The triple combination showed a trend toward synergism in EpS-2 cells, while predominantly additive effects were observed in EpS-1 cells (**Fig. 6G and 6H**), supporting the hypothesis that HDAC inhibition may overcome, at least partially, intrinsic resistance to EZH2-targeted therapies. To assess whether the potentiating effect of HDAC inhibition extends beyond EpS, the triple combination was evaluated in another INI1-deficient malignant rhabdoid tumor (MRT) cell line. MRT shares with EpS the defining molecular event of INI1 loss and the consequent epigenetic dependency on EZH2, making it a biologically relevant model to probe the generalizability of our findings across INI1-deficient tumors (58,59). Consistent with the results observed in EpS-2, SAHA enhanced the cytotoxic activity of TZM and DOX, both as single agents and in combination (Supplementary Fig. S7A). The addition of SAHA to TZM-based treatments resulted in a marked reduction of H3K27me^3^ with a concomitant increase in H3K27ac, accompanied by apoptosis activation as indicated by cleavage of caspase-3 (Supplementary Fig. S7B, Supplementary Table S7). Of note, the triple combination showed a tendency toward synergism (Supplementary Fig. S7C), supporting the hypothesis that HDAC inhibition may broadly potentiate EZH2 inhibition-based therapy across INI1-deficient tumor types, independently of histological context.

**Figure 6.**
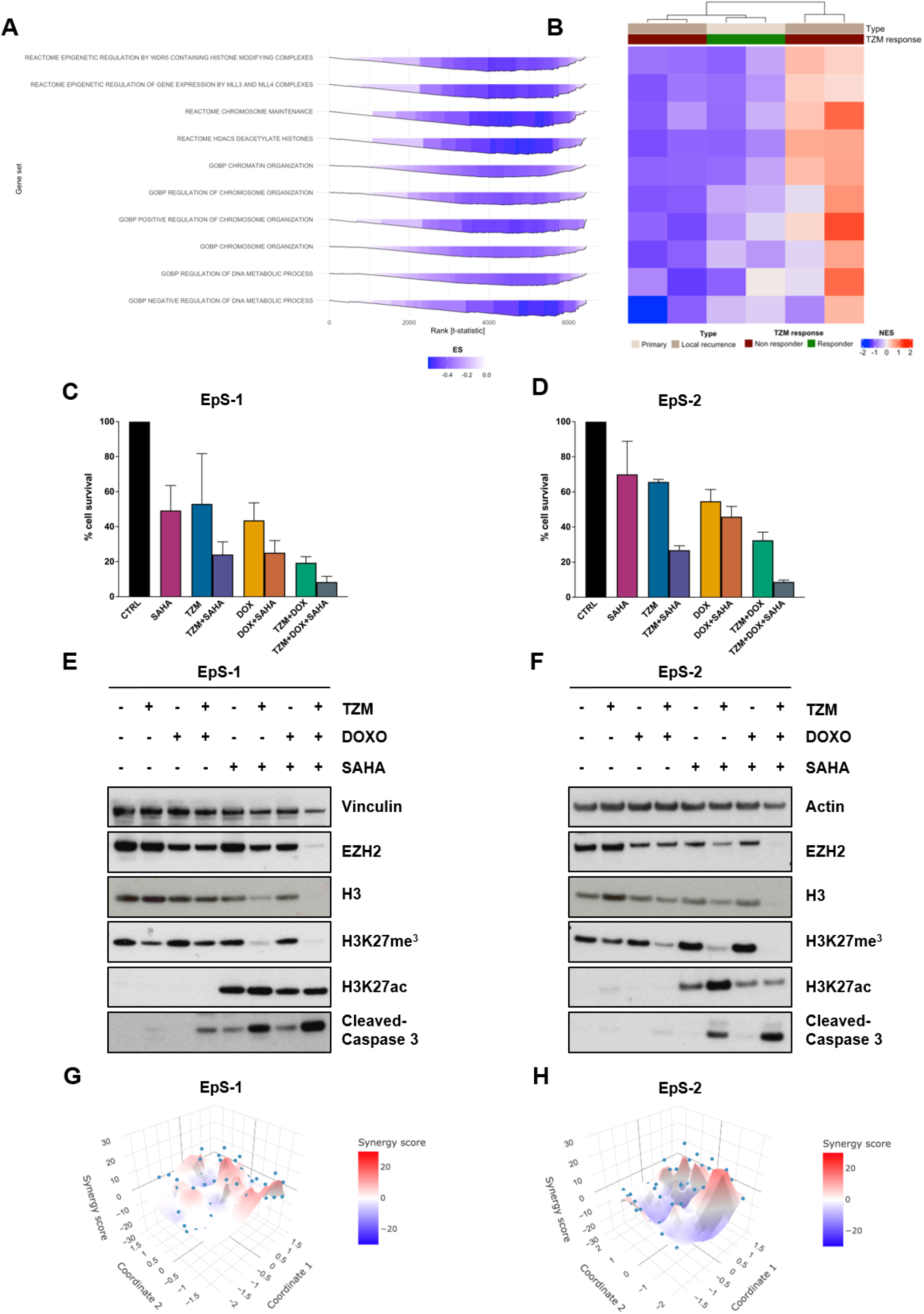
Histone deacetylase (HDAC) activity constrained response to EZH2 inhibition–based therapy in the EpS-2 model. **A,** Gene set enrichment analysis (GSEA) indicating significant downregulation of chromatin structure- and epigenetic regulation-related gene sets in the EpS-1 patient-derived xenograft (PDX) model upon tazemetostat plus doxorubicin (TZM+DOX) treatment. **B**, Heatmap showing hierarchical clustering of single-sample GSEA scores for pathways in A; euclidean metric and ward.D2 linkage measures were used to cluster samples from EpS patients treated with TZM in the EZH-202 phase II trial. Color intensity indicates normalized enrichment scores (NES). **C-D,** Cell survival of EpS-1 **(C)** and EpS-2 **(D)** cells following treatment with TZM, DOX, or their combination in the presence or absence of SAHA. Data are expressed as percentage of viable cells relative to untreated controls. **E-F,** Western blot analysis of chromatin-related markers (EZH2, histone H3, H3K27me³, H3K27ac) and apoptotic activation (caspase-3 cleavage) after treatment with TZM, DOX, or TZM+DOX with or without SAHA in EpS-1 **(E)** and EpS-2 **(F)** cells. Cropped images of selected proteins are shown. Actin and vinculin served as loading controls. **G-H,** Synergy plots for combination treatment with TZM+DOX+SAHA for EpS-1 **(G)** and EpS-2 **(H)** cells. Cells were treated for 72 hours and analyzed for synergy using the ZIP model.

## Discussion

In this study, we identified chromatin remodeling dynamics as a critical determinant of response to EZH2 inhibition-based therapies in EpS. Our data indicate that the therapeutic activity of EZH2 inhibition relies not only on the reduction of the repressive H3K27me3 mark, but also on the subsequent establishment of a more permissive chromatin state characterized by H3K27 acetylation. Failure to induce this epigenetic transition, potentially sustained by increased HDAC activity, limits the efficacy of EZH2-targeted therapy, even when combined with DOX. Importantly, pharmacological HDAC inhibition restores H3K27ac accumulation and sensitizes to EZH2 inhibition-based treatments, providing a rationale for chromatin-directed combination strategies in INI1-deficient malignancies.

Here, we combined EZH2i with DOX in two in-house generated PDX models of P-EpS and their matched cell lines, interrogating both anti-tumor activity and the molecular mechanisms of response and resistance to these treatments. EZH2i and DOX interacted synergistically in EpS-1 *in vitro* and elicited robust antitumor activity *in vivo*, achieving remarkable growth delay in all treated animals (mTVI: 92%). Rechallenge therapy with EZH2i+DOX in the EpS-1/R relapsed PDX model, developed from regrowing EpS-1 treated with the same combination, resulted in a significant tumor volume growth inhibition, suggesting preservation of at least partial sensitivity to rechallenge therapy. Conversely, the EpS-2 displayed intrinsic resistance, with no tumor growth inhibition observed after individual or combined treatments. Together, these models recapitulate two clinically relevant patterns of inadequate response to EZH2-targeted therapy: adaptive persistence despite initial sensitivity (EpS-1) and primary refractoriness (EpS-2). Conversely to what was observed in EpS-1, cytoprotective autophagy contributed to the limited apoptotic response to EZH2i-based regimens in EpS-2, as demonstrated by restoration of apoptosis upon BafA1 co-treatment. Baseline transcriptomic profiling revealed that EpS-2 displayed a transcriptional state enriched for EMT-like programs, confirmed at the protein level by higher vimentin expression, and functionally by greater migratory and invasive capacity than EpS-1. Post-treatment transcriptomic analysis showed that EZH2i-DOX specifically downregulated gene sets linked to chromatin organization, DNA metabolic processes, and HDAC-associated pathways exclusively in EpS-1, with no significant reprogramming detected in EpS-2. Consistently, H3K27ac, a mark of transcriptional activation following EZH2 inhibition, accumulated in EpS-1 but not EpS-2, leading to the hypothesis that HDAC activity could impair EZH2i-induced chromatin remodeling. Addition of the pan-HDAC inhibitor SAHA to the EZH2i-DOX combination increased H3K27ac levels in EpS-2 and the MRT G-401 cell lines, induced apoptosis, and showed a tendency toward synergism. These findings suggest HDAC inhibition as a rational strategy to overcome resistance to EZH2i-based therapy not only in EpS, but also in other INI1-deficient tumors, such as MRT. The inclusion of MRT models was based on their shared loss of INI1 expression and consequent epigenetic dependency on EZH2, making them a biologically relevant system to assess whether the effects of HDAC inhibition could extend beyond EpS.

Our study carries some limitations. First, our experimental platform comprises only two PDX models with their corresponding cell lines, and the relapsed EpS-1/R model derived from regrowing EpS-1 tumors after EZH2i-DOX treatment cessation. Although these models were previously comprehensively characterized and validated as representative of human P-EpS, recapitulating key histomorphological and molecular features and clustering within the landscape of clinical EpS samples based on both DNA methylation and transcriptomic profiles (13), they are unlikely to capture the full spectrum of biological heterogeneity of this disease. EpS, even when restricted to the proximal-type, encompasses significant interpatient variability in genomic complexity and epigenetic landscape (13,60,61), as exemplified by the markedly different chromosomal architectures of EpS-1 and EpS-2. This degree of molecular divergence, while biologically meaningful and reflective of the heterogeneity that exists even within a single EpS subtype, limits our ability to reduce the observed differences in drug response to a single molecular biomarker or a gene expression signature. The same consideration applies to the enhanced EMT-like features identified in EpS-2. Although baseline enrichment of EMT-related pathways was associated with intrinsic resistance, our data do not establish whether these programs actively drive resistance or rather reflect a broader transcriptional cell state associated with reduced susceptibility to EZH2 inhibition-based therapy. Future studies incorporating additional P-EpS models, single-cell transcriptomic profiling, and longitudinal analyses of matched parental and regrowing EpS-1 tumors will be instrumental to distinguish causal mechanisms from correlative cell-state signatures. In this context, Kazansky et al. identified prospective predictors of EZH2i response, including alterations of the RB1/E2F transcriptional axis and overexpression of PRICKLE1, by analyzing tumors collected from a cohort of patients enrolled in a phase II clinical trial of TZM (57,62). Additionally, both PDX models belong to the proximal subtype of EpS. Whether the findings described here apply to the distal variant, which carries a more favorable prognosis and distinct clinical behavior, remains to be investigated (35).

Our study provides the biological basis for additional preclinical investigation aimed at developing rational epigenetic combination strategies for EpS. Loss of INI1 expression shifts the balance between PRC2-driven repression and SWI/SNF-mediated activation toward aberrant chromatin compaction (63), rendering EZH2, the principal driver of H3K27me^3^, a therapeutically actionable vulnerability in EpS and other INI1-deficient malignancies, such as MRT (64). However, based on our data, EZH2 inhibition alone is insufficient to collapse this epigenetic dependency when compensatory mechanisms, such as HDAC-mediated maintenance of a hypoacetylated H3K27 chromatin state, remain operative. In EpS specifically, the therapeutic relevance of this crosstalk was first explored by Lopez et al., who demonstrated that pan-HDAC inhibition with abexinostat exerts potent antiproliferative activity in three EpS cell lines and in xenograft models, reducing EZH2 expression and inducing G2/M cell cycle arrest and apoptosis (65). Our study extends this observation by providing functional evidence that HDAC activity mechanistically limits H3K27ac accumulation in response to EZH2i in EpS, and that pharmacological HDAC inhibition can restore this response and sensitize an intrinsically resistant model to EZH2-targeted therapy. Although the present study employed the pan-HDAC inhibitor SAHA and therefore does not allow attribution of the observed effects to specific HDAC isoforms, our findings raise the possibility that class I HDACs contribute to maintaining H3K27 in a hypoacetylated state, thereby limiting the chromatin-opening effects of EZH2i. Future studies using class I-selective HDAC inhibitors or genetic perturbation approaches will be required to determine which HDAC family members mediate this phenotype. Notably, the evidence that SAHA similarly potentiates EZH2i-based therapy in MRT suggests that HDAC-mediated suppression of H3K27ac may represent a shared resistance mechanism across the broader family of INI1-deficient malignancies.

Several groups have recently mapped additional targetable vulnerabilities emerging downstream of EZH2 inhibition. Kazansky et al. demonstrated that acquired resistance to EZH2i in EpS and MRT models is driven by activation of the RB1/E2F transcriptional axis and that targeting Aurora kinase B (AURKB) can circumvent this resistance (57). Building on these findings, they subsequently demonstrated that EZH2 inhibition itself generates a *bona fide* synthetic lethal vulnerability. Specifically, by upregulating the transposase-derived nuclease PGBD5, TZM promotes endogenous DNA damage in EpS and MRT cells, creating an induced dependency on ATR-mediated DNA repair that can be pharmacologically exploited by the ATR inhibitor elimusertib (62). Collectively, these findings show that INI1 loss generates a multilayered dependency that can be targeted through distinct but complementary strategies in EpS and, more broadly, in INI1-deficient tumors. While previously described approaches exploit vulnerabilities that emerge downstream of EZH2 inhibition, our findings position HDAC inhibition as a more proximal strategy acting directly at the level of chromatin regulation. Specifically, HDAC inhibition enhances and stabilizes the chromatin-opening effects induced by EZH2i, rather than targeting secondary stress responses that arise in parallel. This mechanistic complementarity is supported by consistent efficacy observed across both EpS and MRT cellular models. The translational relevance of EZH2-HDAC co-targeting within the sarcoma field is further supported by a recent study by Omran et al. in uterine sarcoma cells, demonstrating that the combination of EZH2i with the class I-selective HDAC inhibitor entinostat produces superior antiproliferative and pro-apoptotic effects compared to either single agent in both 2D and 3D cultures, with concomitant reduction of Ki67 and PCNA and upregulation of BAX and activated caspase-3 (66). Beyond the sarcoma context, analogous synergistic relationships between EZH2 and HDAC inhibitors have been described in multiple tumor types, such as castration-resistant prostate cancer and T-cell lymphoma (67,68). These convergent findings across tumor types position HDAC inhibition as a broadly applicable strategy to enhance chromatin accessibility at H3K27 and potentiate EZH2-targeted therapy. Our data extend this concept for the first time to EpS and MRT-specific contexts.

The present findings carry potential translational relevance in a disease where most patients derive limited benefit from available therapies. In this context, the recent withdrawal of EZH2i from all markets by Ipsen in March 2026 must be acknowledged. This decision was driven by safety signals emerging from the confirmatory SYMPHONY-1 trial evaluating EZH2i in combination with lenalidomide and rituximab in follicular lymphoma, a context both therapeutically and biologically distinct from EpS (https://www.ipsen.com/press-release/ipsen-voluntarily-withdraws-tazverik-tazemetostat-in-follicular-lymphoma-and-epithelioid-sarcoma-3251503/). While this decision has immediate implications for clinical availability of the drug, it does not undermine the pathophysiological role of EZH2 in INI1-deficient tumors, and, consequently, the preclinical effort to provide the rationale for new EZH2-targeted combination strategies in EpS. The favorable tolerability documented for TZM in the EZH-202 trial, without cardiotoxicity, low rate of grade 3-4 adverse events, and no treatment-related deaths, makes it well suited as a maintenance backbone, in contrast with DOX, whose cumulative cardiotoxicity limits the number of administrable cycles (69). From a combinatorial standpoint, HDAC inhibitors represent a clinically tractable drug class with an established tolerability profile: SAHA, romidepsin, belinostat, and panobinostat have received FDA approval for the treatment of T-cell lymphomas and multiple myeloma, and several agents have been extensively investigated in early-phase trials across solid tumor histotypes (70). Beyond HDAC inhibition, the toxicity profile of DOX warrants exploration of alternative combination partners for EZH2i in EpS (29). Gemcitabine-based regimens and vinorelbine have shown activity in this histotype in the second-line setting (6,21), and their combination with EZH2i represents a potentially less toxic alternative worthy of preclinical and clinical investigation. Additionally, given the capacity of EZH2 inhibition to remodel the tumor immune microenvironment (71), combinations with immune checkpoint inhibitors represent another avenue of interest, currently under evaluation in a phase I/II clinical trial (ClinicalTrials.gov Identifier: <u>NCT05407441</u>) (72,73). A critical unmet need remains the identification of reliable predictive biomarkers of response and resistance to EZH2i-based therapy. H3K27ac accumulation upon EZH2 inhibition represents a biologically rational pharmacodynamic correlate, but its clinical assessment presents significant methodological challenges that will require dedicated solutions before it can be incorporated into prospective study designs.

In conclusion, this study provides preclinical evidence that EZH2i+DOX may yield meaningful antitumor activity in P-EpS. Additionally, persistence of minimal residual disease, which drives relapse upon treatment withdrawal, and primary refractoriness, rooted in an epigenetic state that resists EZH2i-induced chromatin remodeling, were identified as distinct resistance mechanisms in EpS. The retained sensitivity of EpS-1/R to EZH2i+DOX rechallenge supports the potential role of this combination in both induction and re-induction settings, considering toxicity-related limitations with DOX dose and schedule. Failure to accumulate H3K27ac emerges as a hallmark of resistance to EZH2 inhibition, and pharmacological HDAC blockade with SAHA restores this chromatin transition and sensitizes the intrinsically resistant EpS-2 model to EZH2i-based therapy.

## Acknowledgements

The authors thank Monica Tortoreto for her contribution to this study. This study was supported by the 5×1000 funds (2019 Italian Ministry of Health, financial support for health care research); by the institutional grant BRI 2021 (to A. M. Frezza and S. Pasquali); by the International Accelerator Award funded by Associazione Italiana per la Ricerca sul Cancro [ID #24297]/Cancer Research UK [C56167/A29363]/Fundacion Científica, Asociacion Espanola Contra el Cancer [Foundation AECC-GEACC19007MA] (to P. Huang, A. Gronchi, N. Zaffaroni); and by “Ricerca Corrente” funds from the Italian Ministry of Health. C. Soffientini and S. Pasquali are supported by AIRC Individual Grant–Next Gen Clinician Scientist “Fondazione 13 Marzo”.

## Author contributions

**Conceptualization:** N. Arrighetti, C. Soffientini, V. Zuco, S. Percio, A. Kentsis, P. H. Huang, A. Gronchi, A. M. Frezza, S. Stacchiotti, N. Zaffaroni, S. Pasquali; **Methodology:** N. Zaffaroni, S. Pasquali; **Software:** S. Percio, E. Del Savio, L. Sigalotti, R. Maestro;; **Validation:** N. Arrighetti, C. Soffientini; **Formal analysis:** N. Arrighetti, C. Soffientini, S. Percio, E. Del Savio, L. Sigalotti, R. Maestro, G. P. Dagrada, M. Barisella, P. Collini; **Investigation:** N. Arrighetti, C. Soffientini, V. Zuco, L. Cleris, S. A. Ahmed, E. Del Savio, L. Sigalotti, R. Maestro, S. Brich, G. P. Dagrada, M. Barisella, P. Collini; **Resources:** E. Del Savio, L. Sigalotti, R. Maestro, A. Gronchi, N. Zaffaroni, S. Pasquali; **Data curation:** N. Arrighetti, C. Soffientini, V. Zuco, S. Percio, E. Del Savio, L. Sigalotti, R. Maestro; **Writing – original draft:** N. Arrighetti, C. Soffientini, N. Zaffaroni, S. Pasquali; **Writing – review & editing:** All authors; **Visualization:** N. Arrighetti, C. Soffientini, S. Percio, E. Del Savio, L. Sigalotti, R. Maestro, G. P. Dagrada; **Supervision:** A. Gronchi, A. M. Frezza, S. Stacchiotti, N. Zaffaroni, S. Pasquali; **Project administration:** S. Stacchiotti, N. Zaffaroni, S. Pasquali; **Funding acquisition:** P. H. Huang, A. Gronchi, A. M. Frezza, N. Zaffaroni, S. Pasquali.

