## Supplementary Materials, Figures, and Tables for "Histone deacetylase activity limits the response to EZH2 inhibition-based therapy in epithelioid sarcoma and is targetable by epigenetic combination"

### Supplementary – Summary

### Supplementary Materials

**Development of patient-derived xenograft (PDX) models**

Tumor samples were aseptically dissected and cut into approximately 3 $mm^3$ fragments. At least three fragments were implanted subcutaneously into the right flank of 6–8-week-old female CB17/Icr-Prkdc$^{scid}$/IcrIcoCrl mice (SCID; Charles River Laboratories; RRID: IMSR_CRL:236). Tumor growth was monitored biweekly by measuring two perpendicular diameters with a Vernier caliper, and tumor volume (TV) was calculated according to the formula TV = $d^2 \times D / 2$, where $d$ and $D$ represent the shortest and longest diameters, respectively. When TV reached approximately 600 $mm^3$, tumors were harvested and transplanted into recipient mice. After the third *in vivo* passage, PDXs were considered established.

SCID mice were housed in a pathogen-free facility under controlled temperature and humidity, with free access to food and water. Body weight and clinical conditions were routinely monitored throughout the experiments.

**Caspase-3 catalytic activity assay**

Caspase-3 catalytic activity was measured using the APOPCYTO Caspase-3 Colorimetric Assay Kit (MBL International Corporation, RRID: SCR_013389). Cells were treated with TZM, DOX, or their combination at $IC_{50}$ concentrations for 72 or 144 hours. Following treatment, both floating and adherent cells were collected, lysed, and protein extracts were incubated with the substrate N-acetylAsp-Glu-Val-Asp-AMC (DEVD-AMC). Substrate cleavage was quantified by spectrofluorometry analysis (excitation 380nm, emission 460nm) using POLARstar OPTIMA plate reader (BMG Labtech).

### Supplementary Figures

#### Supplementary Figure 1

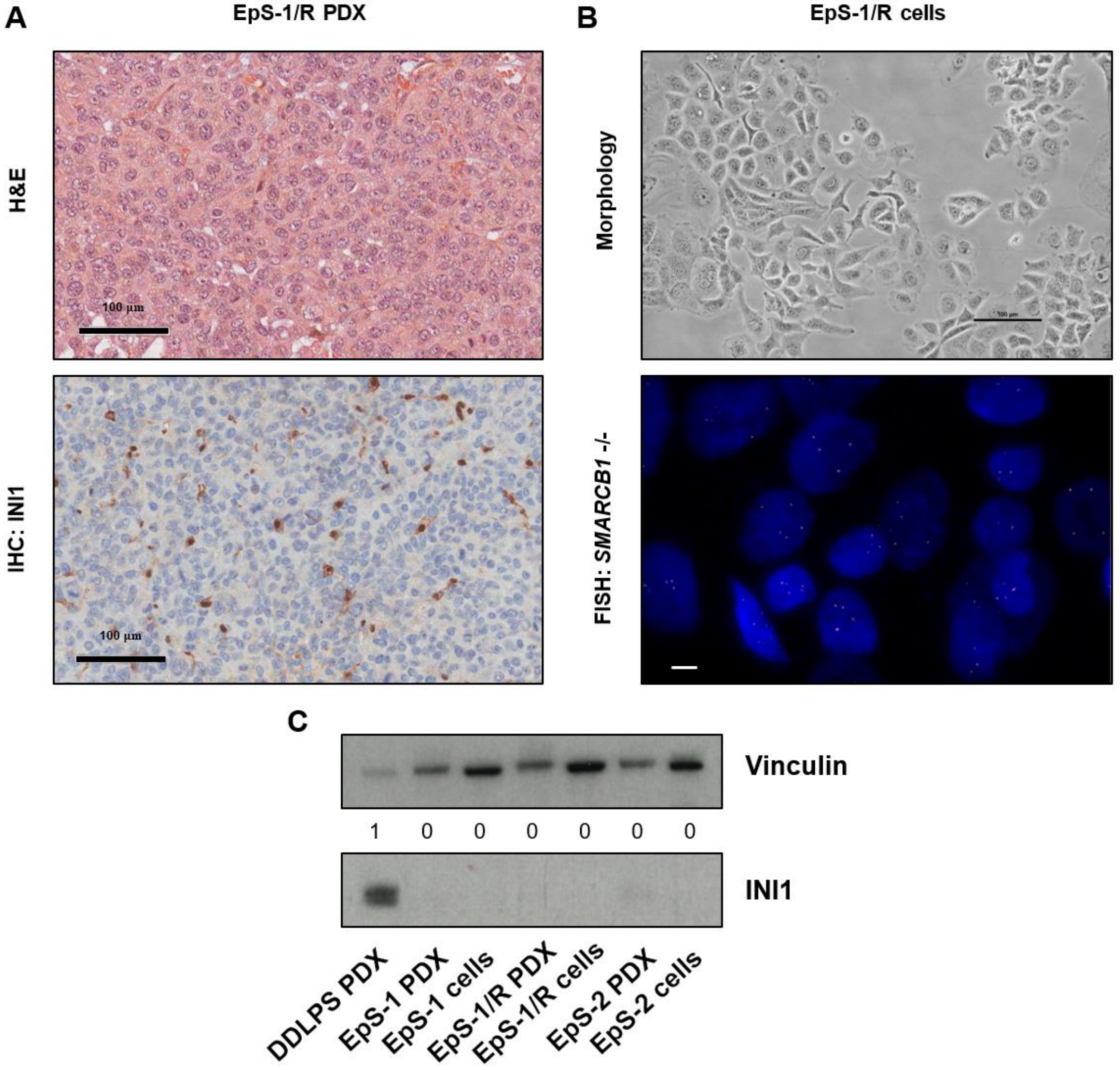

**Supplementary Figure 1. Characterization of the EpS-1/R patient-derived xenograft (PDX) model and matched cell line. A,** Histopathological evaluation of the EpS-1/R PDX by haematoxylin and eosin (H&E) staining (upper panel; scale bar, 100 µm). INI1 protein expression was assessed by immunohistochemistry (IHC) (lower panel; scale bar, 100 µm). **B,** Morphology of EpS-1/R cells derived from the corresponding PDX model, growing as monolayer (upper panel; scale bars, 100 µm). FISH analysis using the in-house dual-color probe (SMARCB1/PDGFB) showed complete loss of *SMARCB1* signals in neoplastic cells (lower panel; scale bars, 10 µm). **C,** Western blot analysis with band intensities quantification showing absence of INI1 protein expression in EpS-1, EpS-1/R, and EpS-2 PDX models and in their corresponding cell lines. Representative cropped images are shown. A dedifferentiated liposarcoma (DDLPS) PDX was used as positive control. Vinculin served as loading control.

#### Supplementary Figure 2

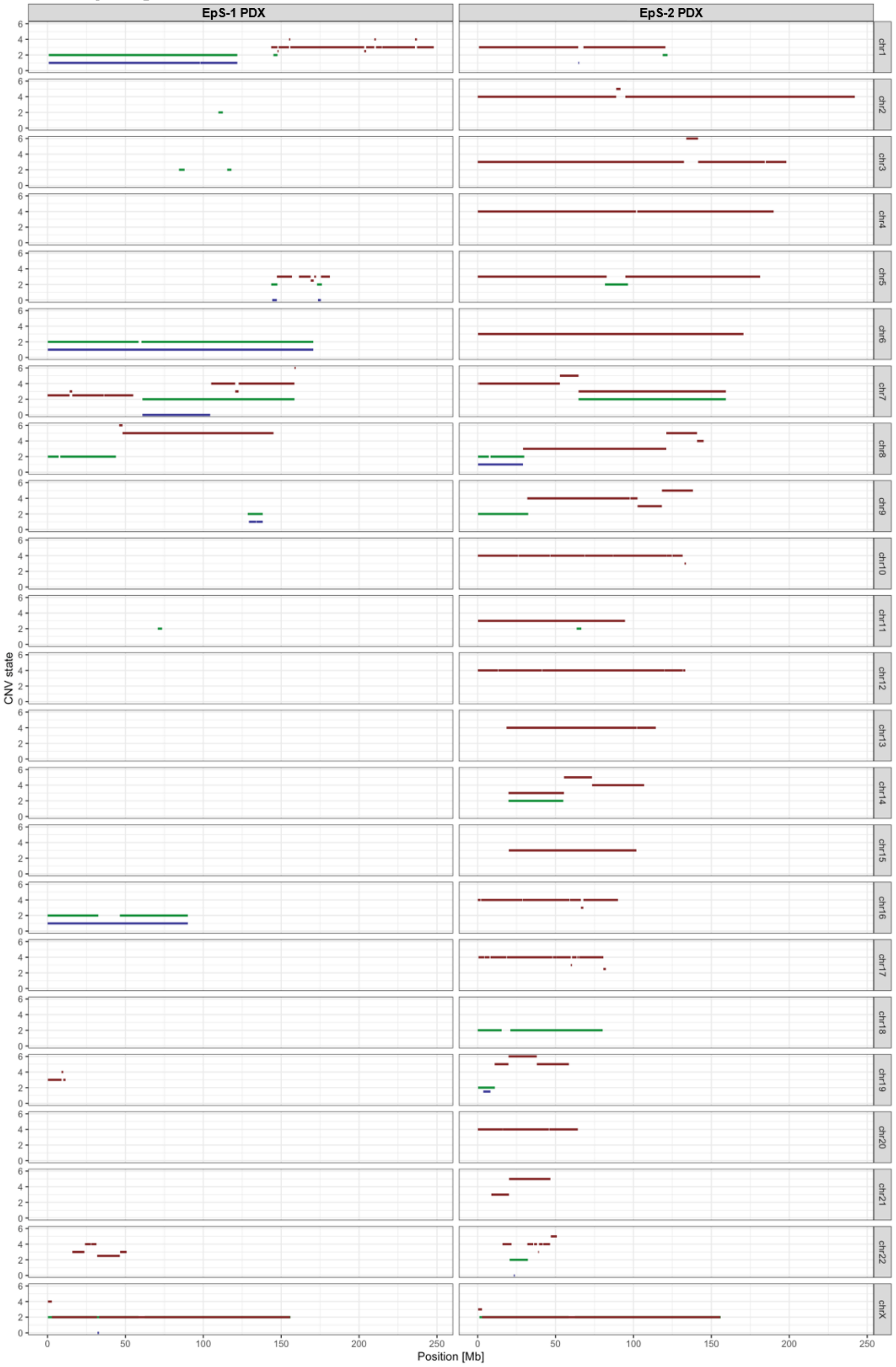

**Supplementary Figure 2. Genome-wide copy number profiles of EpS-1 and EpS-2 patient-derived xenograft (PDX) models at baseline.** Segment plots showing copy number variation (CNV) states across all autosomes and the X chromosome for EpS-1 and EpS-2 PDX models at baseline, as determined by the OncoScan™ CNV Plus Assay (Thermo Fisher Scientific Inc.). The x-axis represents chromosomal position (Mb) and the y-axis indicates the CNV state, with gains (red), losses (blue), and loss of heterozygosity (LOH) (green) shown per segment.

#### Supplementary Figure 3

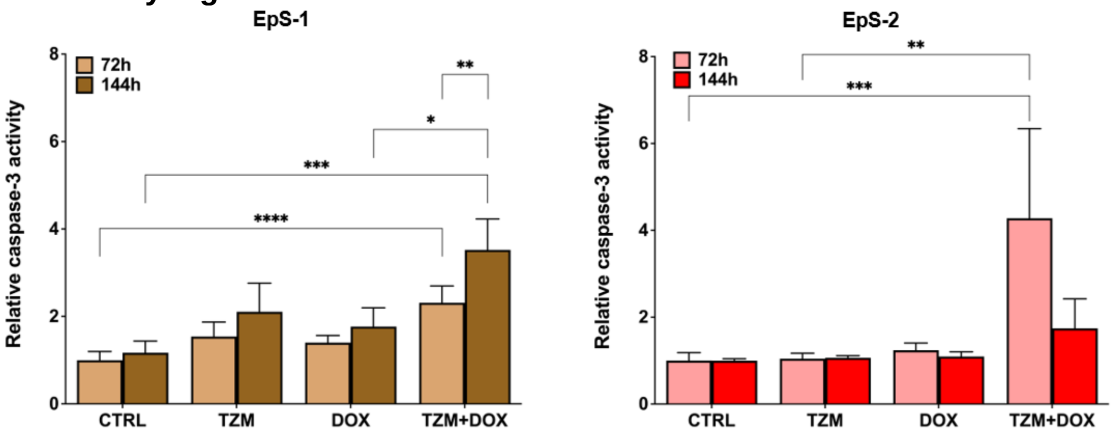

**Supplementary Figure 3. Caspase-3 enzymatic activity following treatment with tazemetostat (TZM) and doxorubicin (DOX), alone or in combination, in EpS-1 and EpS-2 cell lines.** Caspase-3 activity was assessed after 72 and 144 hours of treatment with TZM and DOX, administered as single agents or in combination, in EpS-1 and EpS-2 cells using a commercial enzymatic reaction reagent kit. Data represent mean ± SD of three independent experiments. Statistical analysis: comparisons among treatment groups at each time point were performed using the Kruskal–Wallis test followed by Dunn's multiple comparison test. The comparison between 72 and 144 hours within the TZM+DOX group was performed using the Mann–Whitney test. *p < 0.05; **p < 0.01; ***p < 0.001; ****p < 0.0001.

#### Supplementary Figure 4

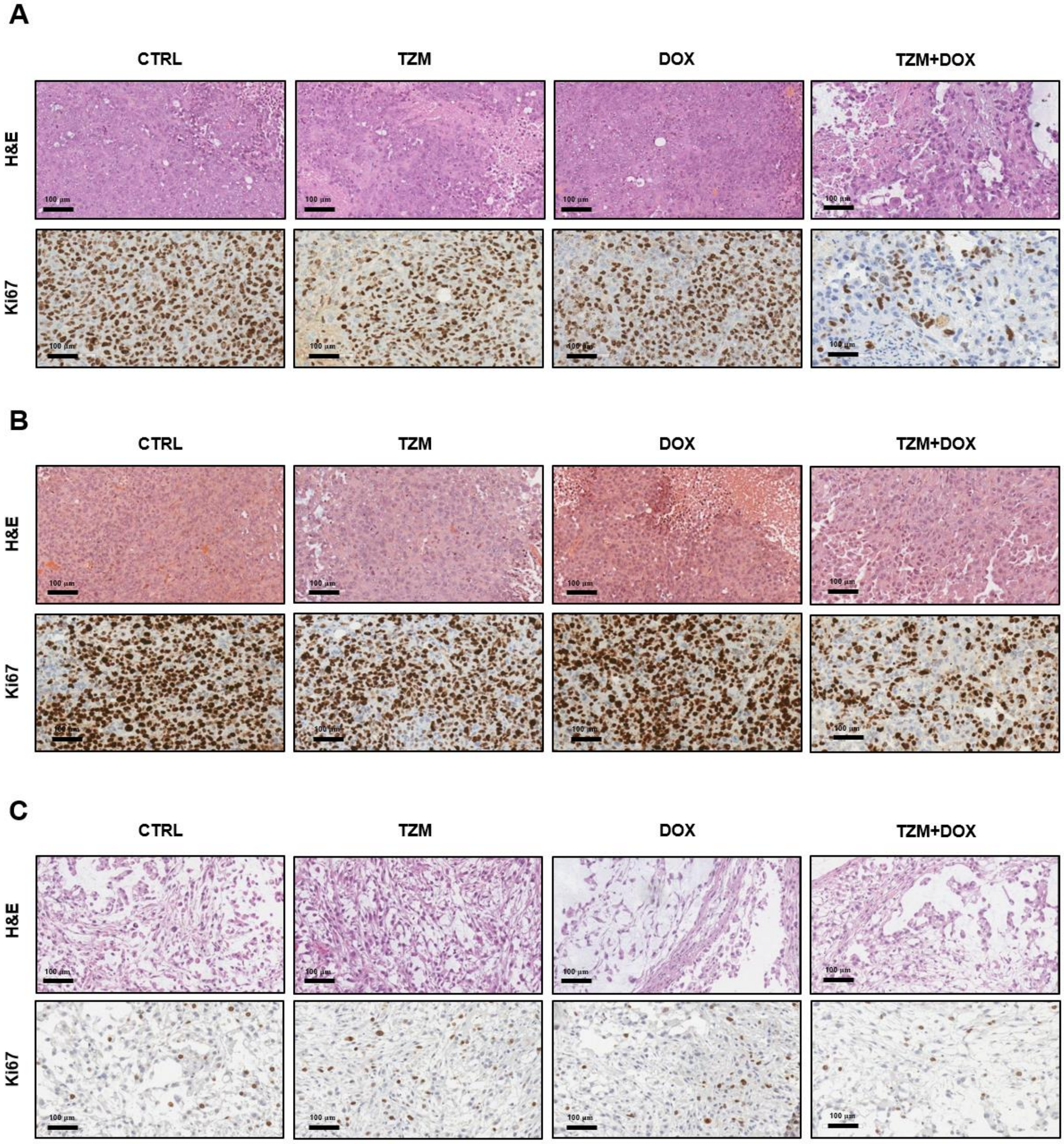

**Supplementary Figure 4. In vivo activity of tazemetostat (TZM) and doxorubicin (DOX), alone or in combination. A,** Histomorphological evaluation (upper panels) and Ki67 immunohistochemical staining (lower panels) of EpS-1 tumors collected from untreated and treated mice at the end of therapy. Scale bar, 100 µm. **B,** Histomorphological evaluation (upper panels) and Ki67 immunohistochemical staining (lower panels) of EpS-1/R tumors collected from untreated and treated mice at the end of therapy. Scale bar, 100 µm. **C,** Histomorphological evaluation (upper panels) and Ki67 immunohistochemical staining (lower panels) of EpS-2 tumors collected from untreated and treated mice at the end of therapy. Scale bar, 100 µm.

#### Supplementary Figure 5

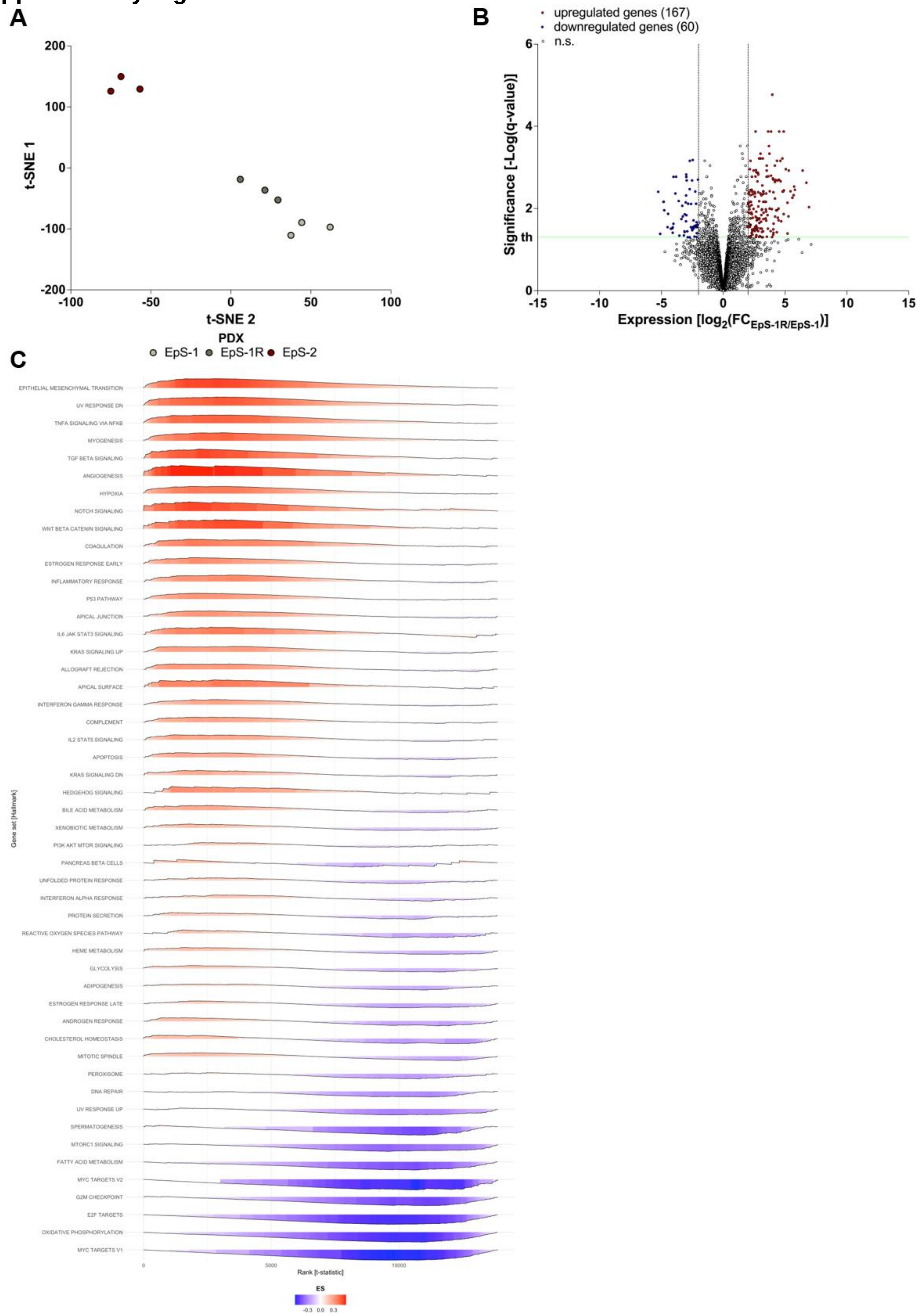

**Supplementary Figure 5. Baseline transcriptomic profiling of EpS-1, EpS-1/R, and EpS-2 patient-derived xenograft (PDX) models at baseline. A,** t-distributed stochastic neighbor embedding (t-SNE) plot of baseline transcriptomic profiles from EpS-1, EpS-1/R, and EpS-2 PDX models. Three independent tumors were sequenced for each PDX model. **B,**

Volcano plot depicting differential gene expression between EpS-1 and EpS-1/R PDX models at baseline. **C,** Ridge plot showing gene set enrichment analysis (GSEA) results for MSigDB Hallmark collection in EpS-2 relative to EpS-1 PDX models. The color of each ridge represents the enrichment score (ES), ranging from red (positive enrichment in EpS-2) to blue (negative enrichment, i.e., enrichment in EpS-1). Gene sets are ordered from the most enriched in EpS-2 to the least enriched.

**Supplementary Figure 6**

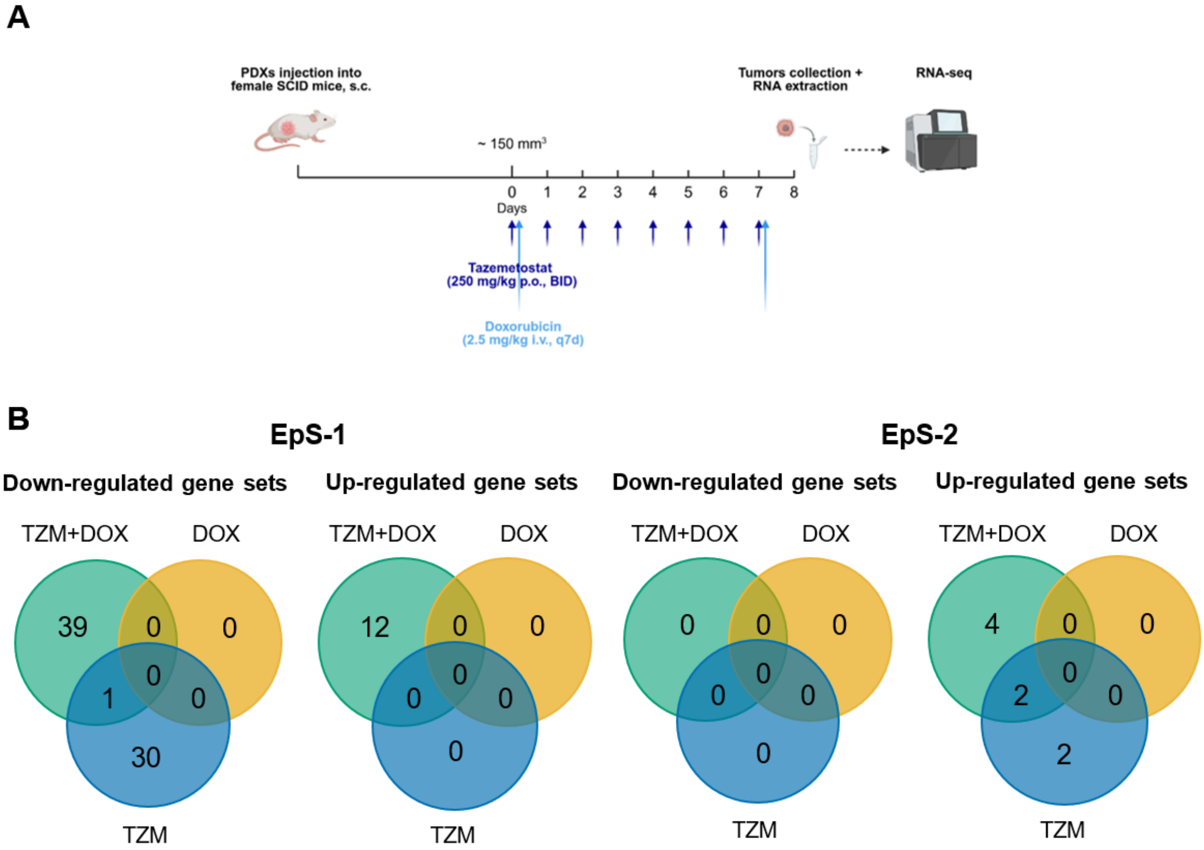

**Supplementary Figure 6. Transcriptional profiling of EpS-1 and EpS-2 patient-derived xenograft (PDX) models following treatment with tazemetostat (TZM) and doxorubicin (DOX), alone or in combination. A,** Schematic representation of the short-term *in vivo* treatment protocol performed in EpS-1 and EpS-2 PDX models for tumor collection and subsequent RNA-sequencing analysis. TZM and DOX were administered as single agents or in combination according to the indicated schedules. **B,** Venn diagrams showing the number of significantly upregulated and downregulated gene sets identified upon drug treatments in EpS-1 and EpS-2 PDX models. Gene set enrichment analysis (GSEA) was performed using the following databases: Hallmark, GO:BP, KEGG Pathway, and Reactome. Statistical significance threshold: adjusted $p < 0.1$.

#### Supplementary Figure 7

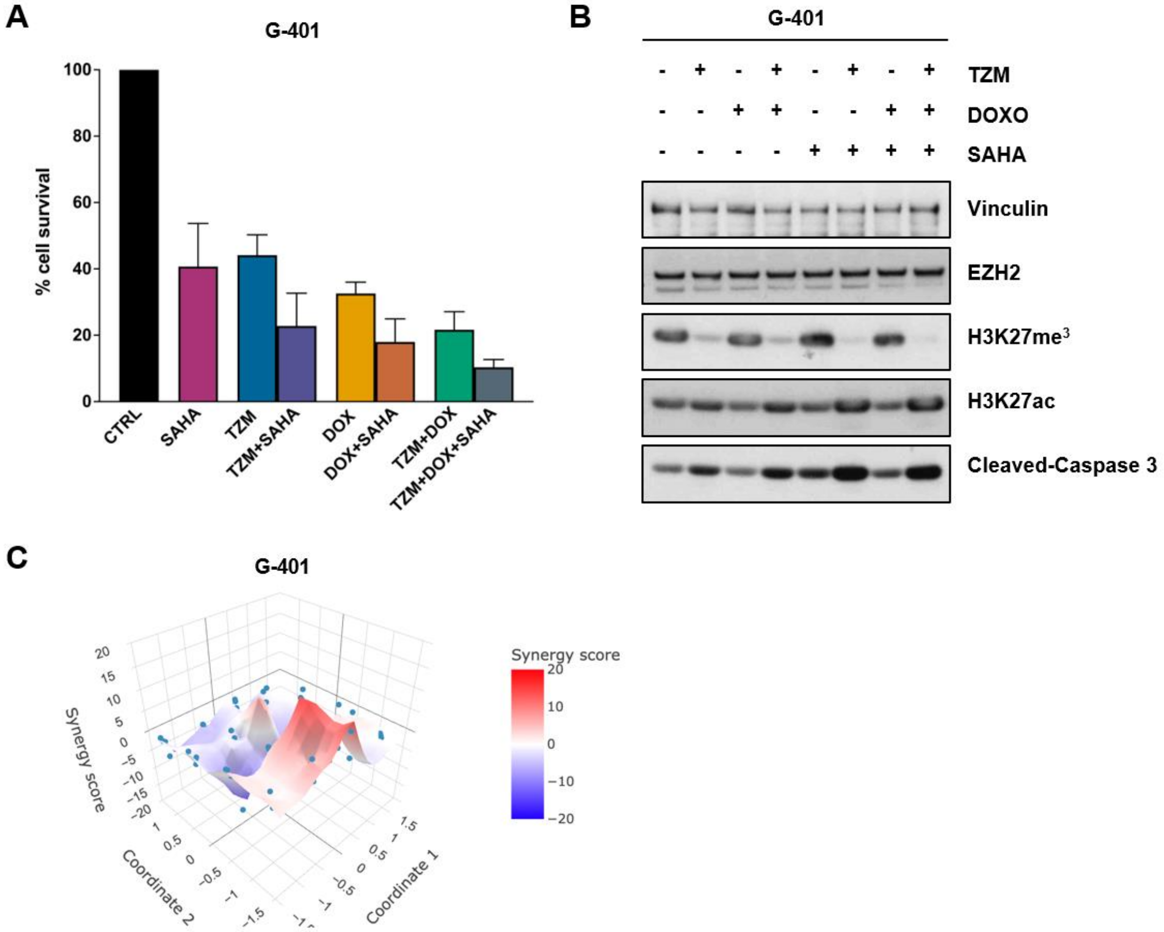

**Supplementary Figure 7. Histone deacetylase (HDAC) activity constrained response to EZH2 inhibition–based therapy in the G-401 cell line. A,** Cell survival of G-401 cells following treatment with tazemetostat (TZM), doxorubicin (DOX), or their combination in the presence or absence of SAHA. Data are expressed as percentage of viable cells relative to untreated controls. **B,** Western blot analysis of chromatin-related markers (EZH2, H3K27me$^3$, H3K27ac) and apoptotic activation (caspase-3 cleavage) after treatment with TZM, DOX, or TZM+DOX with or without SAHA in G-401 cells. Cropped images of selected proteins are shown. Actin and vinculin served as loading controls. **C,** Synergy analysis of the triple combination TZM+DOX+SAHA in G-401 cells.

### Supplementary Tables

#### Supplementary Table 1

| Antibody | Supplier | Catalog number | RRID |
|---|---|---|---|
| Anti-ß-actin | Abcam | ab8226 | AB_306371 |
| Anti-ß-catenin | Cell Signaling Technology | 8480 | AB_11127855 |
| Anti-cleaved-caspase-3 | Cell Signaling Technology | 9661 | AB_2341188 |
| Anti-E-cadherin | Santa Cruz Biotechnology | sc-8426 | AB_626780 |
| Anti-EZH2 | Cell Signaling Technology | 5246 | AB_10694683 |
| Anti-GAPDH | Sigma-Aldrich | G8795 | AB_1078991 |
| Anti-H3 | Thermo Fischer Scientific | PA5-16183 | AB_10985434 |
| Anti-H3K27ac | Cell Signaling Technology | 8173 | AB_10949503 |
| Anti-H3K27me$^3$ | Cell Signaling Technology | 9733 | AB_2616029 |
| Anti-LC3B | Cell Signaling Technology | 83506 | AB_2800018 |
| Anti-PARP | Cell Signaling Technology | 9542 | AB_2160739 |
| Anti-INI1 | Cell Signaling Technology | 91735 | AB_2800172 |
| Anti-ß-tubulin | Sigma-Aldrich | T4026 | AB_477577 |
| Anti-vimentin | Dako | MO725 | AB_10013485 |
| Anti-vinculin | Sigma-Aldrich | V9131 | AB_477629 |
| Anti-rabbit IgG | Cell Signaling Technology | 7074 | AB_2099233 |
| Anti-mouse IgG | Cell Signaling Technology | 7076 | AB_330924 |

**Supplementary Table 1. List of antibodies used for western blotting.** Supplier, Catalog number, and Research Resource Identifier (RRID) are indicated for each antibody.

#### Supplementary Table 2

| EpS-1 | | | | | | | | EpS-2 | | | | | | | | |
|---|---|---|---|---|---|---|---|---|---|---|---|---|---|---|---|---|
| 72h | | | | 144h | | | | 72h | | | | 144h | | | | |
| - | + | - | + | - | + | - | + | - | + | - | + | - | + | - | + | **TZM** |
| - | - | + | + | - | - | + | + | - | - | + | + | - | - | + | + | **DOX** |
| 1.0 | 0.7 | 0.6 | 0.5 | 1.0 | 0.5 | 0.4 | 0.4 | 1.0 | 1.9 | 1.5 | 1.7 | 1.0 | 1.7 | 1.5 | 0.6 | **H3** |
| 1.0 | 0.3 | 1.9 | 0.4 | 1.0 | 0.1 | 2.3 | 0.1 | 1.0 | 0.2 | 0.7 | 0.4 | 1.0 | 0.2 | 0.8 | 0.5 | **H3K27me$^{3}$** |
| 1.0 | 1.5 | 0.3 | 3.9 | 1.0 | 4.8 | 0.5 | 2.5 | 1.0 | 0.7 | 0.7 | 0.7 | 1.0 | 0.7 | 0.4 | 1.1 | **H3K27ac** |
| 1.0 | 9.7 | 0.6 | 44.0 | 1.0 | 7.0 | 0.0 | 5.9 | 1.0 | 9.4 | 16.0 | 1399.2 | 1.0 | 1.7 | 0.9 | 1.1 | **Cleaved Caspase-3** |
| 1.0 | 71.1 | 15.2 | 746.4 | 1.0 | 2.3 | 0.1 | 2.6 | 1.0 | 98.3 | 50.2 | 183.1 | 1.0 | 3.0 | 1.0 | 3.2 | **LC3B-II** |

**Supplementary Table 2. Quantification of band intensities for blot reported in Figure 2C.** Quantification was performed using ImageJ 1.47q software. Total histone H3, cleaved caspase-3, and LC3B-II signals were normalized to the corresponding β-tubulin or vinculin loading control, whereas H3K27me$^{3}$ and H3K27ac signals were normalized to total histone H3. Normalized values for each treatment condition (TZM, DOX, and TZM+DOX) were expressed relative to the untreated control sample.

#### Supplementary Table 3

| EpS-1 | | | | | EpS-2 | | | | | |
|---|---|---|---|---|---|---|---|---|---|---|
| - | - | + | - | + | - | - | + | - | + | **TZM** |
| - | - | - | + | + | - | - | - | + | + | **DOX** |
| - | + | + | + | + | - | + | + | + | + | **BafA1** |
| 1.0 | 563815.1 | 1445057.6 | 876221.6 | 2522798.2 | 1.0 | 630.6 | 1400.1 | 164.4 | 1928.6 | **Cleaved Caspase-3** |
| 1.0 | 297.4 | 293.9 | 262.1 | 266.6 | 1.0 | 174.2 | 57.6 | 65.4 | 138.1 | **LC3B-I** |
| 1.0 | 302.9 | 453.2 | 204.9 | 415.6 | 1.0 | 276.0 | 117.1 | 32.5 | 57.7 | **LC3B-II** |
| 1.0 | 0.4 | 0.2 | 0.2 | 0.0 | 1.0 | 0.8 | 0.6 | 0.5 | 0.2 | **PARP** |
| 1.0 | 121.0 | 128.3 | 82.3 | 87.9 | 1.0 | 729.2 | 1956.0 | 105.5 | 2609.4 | **Cleaved PARP** |

**Supplementary Table 3. Quantification of band intensities for blot reported in Figure 3C.** Quantification was performed using ImageJ 1.47q software. Signals were normalized to the corresponding actin loading control. Normalized values for each treatment condition (TZM, DOX, and TZM+DOX) were expressed relative to the untreated control sample.

#### Supplementary Table 4

| EpS-1 PDX | | | | EpS-1/R PDX | | | | EpS-2 PDX | | | | |
|---|---|---|---|---|---|---|---|---|---|---|---|---|
| - | + | - | + | - | + | - | + | - | + | - | + | **TZM** |
| - | - | + | + | - | - | + | + | - | - | + | + | **DOX** |
| 1.0 | 0.9 | 0.6 | 0.5 | 1.0 | 1.4 | 0.8 | 1.0 | 1.0 | 0.6 | 0.6 | 0.7 | **H3** |
| 1.0 | 0.1 | 1.0 | 0.2 | 1.0 | 0.0 | 1.2 | 0.2 | 1.0 | 0.0 | 0.7 | 0.3 | **H3K27me$^3$** |
| 1.0 | 1.8 | 1.0 | 1.7 | 1.0 | 3.0 | 0.9 | 2.3 | 1.0 | 1.7 | 1.3 | 1.1 | **H3K27ac** |

**Supplementary Table 4. Quantification of band intensities for blot reported in Figure 4K.** Quantification was performed using ImageJ 1.47q software. Total histone H3 signals were normalized to the corresponding β-actin loading control, whereas H3K27me$^3$ and H3K27ac signals were normalized to total histone H3. Normalized values for each treatment condition (TZM, DOX, and TZM+DOX) were expressed relative to the untreated control sample.

#### Supplementary Table 5

| EpS-1 PDX | EpS-2 PDX | |
|---|---|---|
| 1.0 | 0.0 | **B-Catenin** |
| 1.0 | 2.1 | **Vimentin** |
| 1.0 | 0.0 | **E-Cadherin** |

**Supplementary Table 5. Quantification of band intensities for blot reported in Figure 5D.** Quantification was performed using ImageJ 1.47q software. Signals were normalized to the corresponding actin or vinculin loading control. Normalized values for each patient-derived xenograft (PDX) models were expressed relative to the EpS-1 PDX sample.

#### Supplementary Table 6

| EpS-1 | | | | | | | | EpS-2 | | | | | | | | |
|---|---|---|---|---|---|---|---|---|---|---|---|---|---|---|---|---|
| **-** | **+** | **-** | **+** | **-** | **+** | **-** | **+** | **-** | **+** | **-** | **+** | **-** | **+** | **-** | **+** | **TZM** |
| **-** | **-** | **+** | **+** | **-** | **-** | **+** | **+** | **-** | **-** | **+** | **+** | **-** | **-** | **+** | **+** | **DOX** |
| **-** | **-** | **-** | **-** | **+** | **+** | **+** | **+** | **-** | **-** | **-** | **-** | **+** | **+** | **+** | **+** | **SAHA** |
| 1.0 | 1.1 | 0.7 | 0.7 | 1.3 | 0.8 | 0.8 | 0.0 | 1.0 | 0.9 | 0.4 | 0.4 | 0.5 | 0.2 | 0.5 | 0.0 | **EZH2** |
| 1.0 | 1.2 | 0.9 | 0.9 | 0.8 | 0.2 | 0.7 | 0.0 | 1.0 | 1.5 | 0.8 | 0.4 | 0.9 | 0.5 | 0.9 | 0.0 | **H3** |
| 1.0 | 0.4 | 1.3 | 0.9 | 1.4 | 0.7 | 1.4 | 6.2 | 1.0 | 0.4 | 1.1 | 0.4 | 1.5 | 0.4 | 1.7 | 0.4 | **H3K27me$^3$** |
| 1.0 | 0.1 | 1.1 | 3.1 | 551.4 | 2772.9 | 453.5 | 141502.7 | 1.0 | 60.8 | 9.5 | 26.5 | 1847.3 | 8696.9 | 1238.0 | 34505.5 | **H3K27ac** |
| 1.0 | 9.9 | 0.6 | 174.4 | 180.2 | 744.7 | 197.9 | 1748.7 | 1.0 | 2.4 | 0.1 | 5.9 | 1.5 | 150.6 | 5.3 | 392.3 | **Cleaved Caspase-3** |

**Supplementary Table 6. Quantification of band intensities for blot reported in Figure 6E-F.** Quantification was performed using ImageJ 1.47q software. EZH2, total histone H3, and cleaved caspase-3 signals were normalized to the corresponding β-actin or vinculin loading control, whereas H3K27me$^3$ and H3K27ac signals were normalized to total histone H3. Normalized values for each treatment condition (TZM, DOX, TZM+DOX, SAHA, TZM+SAHA, DOX+SAHA, and TZM+DOX+SAHA) were expressed relative to the untreated control sample.

#### Supplementary Table 7

| G-401 | | | | | | | | |
|---|---|---|---|---|---|---|---|---|
| - | + | - | + | - | + | - | + | **TZM** |
| - | - | + | + | - | - | + | + | **DOX** |
| - | - | - | - | + | + | + | + | **SAHA** |
| 1.0 | 1.5 | 1.3 | 2.1 | 2.1 | 2.1 | 1.9 | 1.9 | **EZH2** |
| 1.0 | 0.6 | 1.2 | 0.8 | 3.0 | 0.5 | 1.8 | 0.2 | **H3K27me$^{3}$** |
| 1.0 | 2.1 | 1.2 | 3.4 | 2.6 | 4.2 | 2.2 | 3.7 | **H3K27ac** |
| 1.0 | 4.2 | 2.0 | 7.1 | 7.7 | 11.1 | 4.9 | 8.6 | **Cleaved Caspase-3** |

**Supplementary Table 7. Quantification of band intensities for blot reported in Supplementary Fig. S7B.** Quantification was performed using ImageJ 1.47q software. EZH2, cleaved caspase-3, H3K27me3, and H3K27ac signals were normalized to the corresponding vinculin loading control. Normalized values for each treatment condition (TZM, DOX, TZM+DOX, SAHA, TZM+SAHA, DOX+SAHA, and TZM+DOX+SAHA) were expressed relative to the untreated control sample.
